# CD180 identifies chemoresistant stem-like blasts and reveals a KMT2A-driven vulnerability in acute myeloid leukaemia

**DOI:** 10.64898/2026.04.23.720316

**Authors:** Madonna M Eltoukhy, Alexander Winton, Francesca Fasanella Masci, Elżbieta Kania, Mary T Scott, Alastair L Smith, Evie Rigby, Alexandra Curran, Aya Gouma, Jennifer Cassels, Lijun Liu, Thomas Stevens, Karen Dunn, Kevin M Rattigan, Meaad Almowaled, Lucy Wheeler, G. Vignir Helgason, Anindita Roy, Pamela Kearns, Paul Wetherell, Thomas A Milne, Brenda Gibson, Paresh Vyas, Christine J Harrison, David Vetrie, Karen Keeshan

## Abstract

Relapse and chemoresistance remain major challenges in paediatric acute myeloid leukaemia (PAML), particularly in KMT2A-rearranged (KMT2A-r) subtypes where conventional markers such as CD34 are often absent, complicating measurable residual disease (MRD) detection. Leukaemia stem/regenerating cells (LSC/LRC) drive disease initiation, progression, and relapse, sharing stemness and chemoresistance properties that make them critical therapeutic targets. Using high-dimensional spectral flow cytometry, we identified CD180, a Toll-like receptor-like surface protein, as highly expressed on blasts and stem-like populations in KMT2A-r AML, while near absent on normal haematopoietic stem cells (HSCs). PAML KMT2A-r exhibits an unconventional immunophenotype dominated by CD34^⁻^CD180^⁺^ populations. Integrated single-cell transcriptomics and functional profiling revealed CD180^high^ clusters enriched for quiescence, oxidative phosphorylation, and KMT2A/LSC stemness signatures. CD180^⁺^ cells demonstrated robust leukaemia-initiating capacity in xenograft models and persisted through therapy, re-emerging at relapse with phenotypic plasticity. Epigenomic analysis showed CD180 is a direct transcriptional target of the KMT2A::MLLT3 fusion complex, regulated by intragenic enhancers and downregulated by menin and BET inhibitors. Longitudinal single-cell analysis confirmed persistence and clonal evolution of CD180^⁺^ populations during treatment and relapse, underscoring their mechanistic role in chemoresistance and disease progression. In summary, CD180 marks dynamic, relapse-driving populations in KMT2A-r PAML, persists through therapy, and importantly is near absent on normal HSCs, offering a selective therapeutic window. These findings position CD180 as a clinically actionable biomarker for MRD detection and a compelling therapeutic target for eradicating chemoresistant, stem-like cells in paediatric AML.

**Main Points:**

1. CD180 marks chemoresistant, relapse-driving stem-like blasts in KMT2A-r paediatric AML, overcoming CD34-based MRD limitations.
2. Absent on normal HSCs, CD180 is a KMT2A::MLLT3 target and actionable for MRD, relapse prediction, and CD180-directed therapies.

**Novelty:** This study introduces CD180 as a novel biomarker and therapeutic target in AML, particularly KMT2A-rearranged subtypes where conventional markers are often absent. Unlike MRD strategies focused on bulk blasts, CD180 marks chemoresistant, stem-like populations driving relapse, critical reservoirs poorly defined in paediatric AML. This work fills a major gap in prognostic assessment and therapy by enabling precise detection of relapse-driving cells and offering a selective therapeutic window.

## Introduction

Acute myeloid leukaemia (AML) in children remains a devastating disease, with treatment still reliant on conventional cytotoxic chemotherapy +/- stem cell transplantation. The urgent need for safer, more effective therapies is clear. Targeted therapies for paediatric AML (PAML) are extremely limited. The MyeChild 01 clinical trial (1) recently investigated a dose-finding strategy for an antibody-drug-conjugate (ADC) targeting CD33, a marker expressed on AML blasts and some leukaemia stem cell (LSC) populations, but also present on healthy haematopoietic stem cells (HSCs) and myeloid cells (2). LSCs and leukaemia regenerating cells (LRCs) are the reservoirs of leukaemia capable of driving initiation, progression, and relapse in AML. These therapy-resistant populations persist in a quiescent/dormant state and adapt under chemotherapy-induced stress, enabling recurrence (3, 4). Conceptually and functionally, LSCs are defined as the disease-inititating cells whereas LRCs are the therapy-persisting, relapse-driving cells. Both are critical for curative strategies, but LRCs highlight the plasticity of leukaemia. While any LRCs may be a subset of LSCs, not all LSCs become LRCs. Bulk and single cell transcriptomic and epigenetic studies focused on cell hierarchies have shown primitive/stem-like expression profiles associated with relapse (5–7). Thus, both LSC and LRC populations exhibit stemness programs, metabolic flexibility, and immune evasion, making them critical targets for intervention and a promising strategy to eradicate AML and prevent relapse.

Multiparameter flow cytometry (MFC)-based measurable residual disease (MRD) is widely used in AML, detecting leukaemia-associated immunophenotypes (LAIPs) or abnormal populations via different-from-normal (DfN). MRD positivity is ≥0.1% CD45^⁺^ cells (8). However, LAIP antigens of the bulk AML blasts are not necessarily synonymous with LSC antigens, and combining both approaches improves prognosis (9, 10). Many MRD/LSC/LRC antigens lack known functional specificity or therapeutic tractability due to their expression on healthy tissues. LSC/LRCs, typically CD34+/CD38-(and some CD34-), drive relapse even when MRD is negative and represent the chemoresistant reservoir of the disease (11–13). Consequently, the European LeukemiaNet (ELN) has recommended the inclusion of LSC-MRD markers such as CD34, CD117, CD123, CD45, CD33, CD56, CD38, CD36 and CD45RA for MRD detection (8) for both PAML and adult AML (aAML) (14–18). While some antigens are shared with normal/healthy cells, their aberrant expression in leukaemic populations creates a therapeutic window, enabling targeted therapies such as anti-CD33 ADCs (19).

CD180 has emerged as a promising candidate as an LSC/MRD antigen in AML. It is a transmembrane toll-like receptor (TLR)-like protein, structurally similar to TLR4, sharing extracellular leucine-rich repeat domains and the accessory molecule MD-1, but lacking the intracellular TIR domain required for MyD88-mediated signalling. Its mRNA is expressed on AML LSCs (defined as CD34+CD38-) but not on normal HSCs (20, 21). However, previous studies did not include KMT2A-r AML, a common PAML subtype seen to have a dominant CD34-immunophenotypic population due to epigenetic silencing (22). Proteomic screening of AML bone marrow samples from the TCGA LAML dataset identified CD180 as a surface protein absent from normal CD34+ stem cells (23) and with limited expression in other tissues (based on the Human Protein Atlas). These data suggest the potential for CD180 as a therapeutic target with tolerable “on-target, off-cancer” toxicity.

In prior work, we showed enriched *CD180* mRNA expression in a murine model of PAML as compared to aAML (24). Whilst not absent in our aAML experimental model, its enrichment in the paediatric model suggested it may have important relevance to the pathophysiology and potential therapeutic targeting of PAML. We now show CD180 as a distinct PAML blast and LSC/LRC/MRD antigen enriched in primary human KMT2A-rearranged (KMT2A-r) AML. We show that CD180 persists at relapse and that CD180 expressing cells have LSC/LRC-like features. Importantly, this antigen is near absent on normal human HSCs, and therefore offering a path of selective vulnerability in PAML LSC/LRCs, while sparing healthy tissue.

## Materials and Methods

### Cell Lines, Mice, and Patient Samples

Human AML cell lines were cultured under standard conditions and authenticated. Primary AML samples were obtained with informed consent and institutional ethics approval. Immunodeficient NSG, NRG and NRG-W41 mice were used for xenograft studies to assess leukaemia-initiating capacity of CD180^⁺^ populations.

### Single-cell and Bulk RNA-seq, ATAC-seq, and ChIP-seq

Single-cell RNA-seq was performed on longitudinal patient samples using droplet-based platforms, followed by computational analysis to identify CD180^high^ clusters. Bulk RNA-seq was used for transcriptome profiling of sorted CD180^high^ and CD180^low^ fractions. Chromatin accessibility was assessed by ATAC-seq, and epigenomic regulation of CD180 was interrogated using ChIP-seq for KMT2A::MLLT3 and histone marks. Data integration pipelines combined transcriptomic and epigenomic datasets to define regulatory mechanisms and enhancer activity.

### Additional Methods

Further detailed methods and details on flow cytometry, immunophenotyping, xenograft procedures, computational pipelines, knockout and overexpression, FISH, Drug treatments, viability assays, and statistical analyses are provided in the Supplementary Methods.

## Results

### CD180 enriched on paediatric KMT2A-r AML

We analysed CD180 expression in AML (aAML and PAML). Bloodspot (25) and TARGET datasets showed higher CD180 in intermediate/poor prognostic KMT2A-r (t(11q23)/MLL) and good prognostic CBFB::MYH11 (inv(16)) subtypes versus RUNX1-RUNX1T1 (t(8;21)). CD180 was elevated in AML compared to healthy HSCs (Lin−, CD34+, CD38−, CD90+, CD45RA−) and bone marrow (BM) (**Figure 1A-B**). High CD180 correlated with poorer survival in KMT2A-r PAML and aAML (Fig. 1C-D), but not in CBF-AML (**Figure 1C-D, Figure S1A**). We analysed CD180 surface expression with other LSC/LRC markers in blast cells (lin-CD45^low^) across PAML cytogenetic subtypes and compared with aAML to assess age- and subtype-specific patterns using diagnostic samples (KMT2A-r, non-KMT2A-r, and aAML; see **Table S1** and **Figure S1B**). CD34 expression was found to be significantly lower in KMT2A PAML compared to non-KMT2A PAML and aAML (**Figure 1E**). KMT2A-r PAML blasts showed markedly reduced expression of canonical LSC/MRD markers (<10% for CD34, CD123, CD117, CD45RA, CD244) compared to non-KMT2A PAML/aAML, while CD33 remained consistent across groups. In contrast, CD180 was significantly higher in KMT2A-r PAML (80% samples >60% CD180+; mean 69.1% ±23.1%, p<0.0001), with strong correlation to blast burden (**Figure 1F-L**). Other markers (CD56, CD93, CD36) trended higher in PAML (**Figure S1C-E**). Elevated CD180 expression and low levels of other antigens explained minimal correlation with conventional markers (**Figure S1F**). LSC/LRCs occupy immunophenotypic compartments resembling normal haematopoiesis, including GMPs, LMPPs, and CD34-populations (3, 9, 11). KMT2A-r PAML showed reduced subsets (HSCs, MPPs, CMPs, GMPs, MEPs) but increased CD34-CD117+ cells versus non-KMT2A PAML and healthy BM. Despite the underrepresentation of CD34+ subpopulations in KMT2A-r PAML, CD180 was significantly overexpressed across all CD34+ subsets compared to normal controls as well as the CD34-CD117+ compartment (**Figure 1M**) distinguishing KMT2A-r PAML from other AML subtypes and conventional LSC markers.

**Figure 1:**
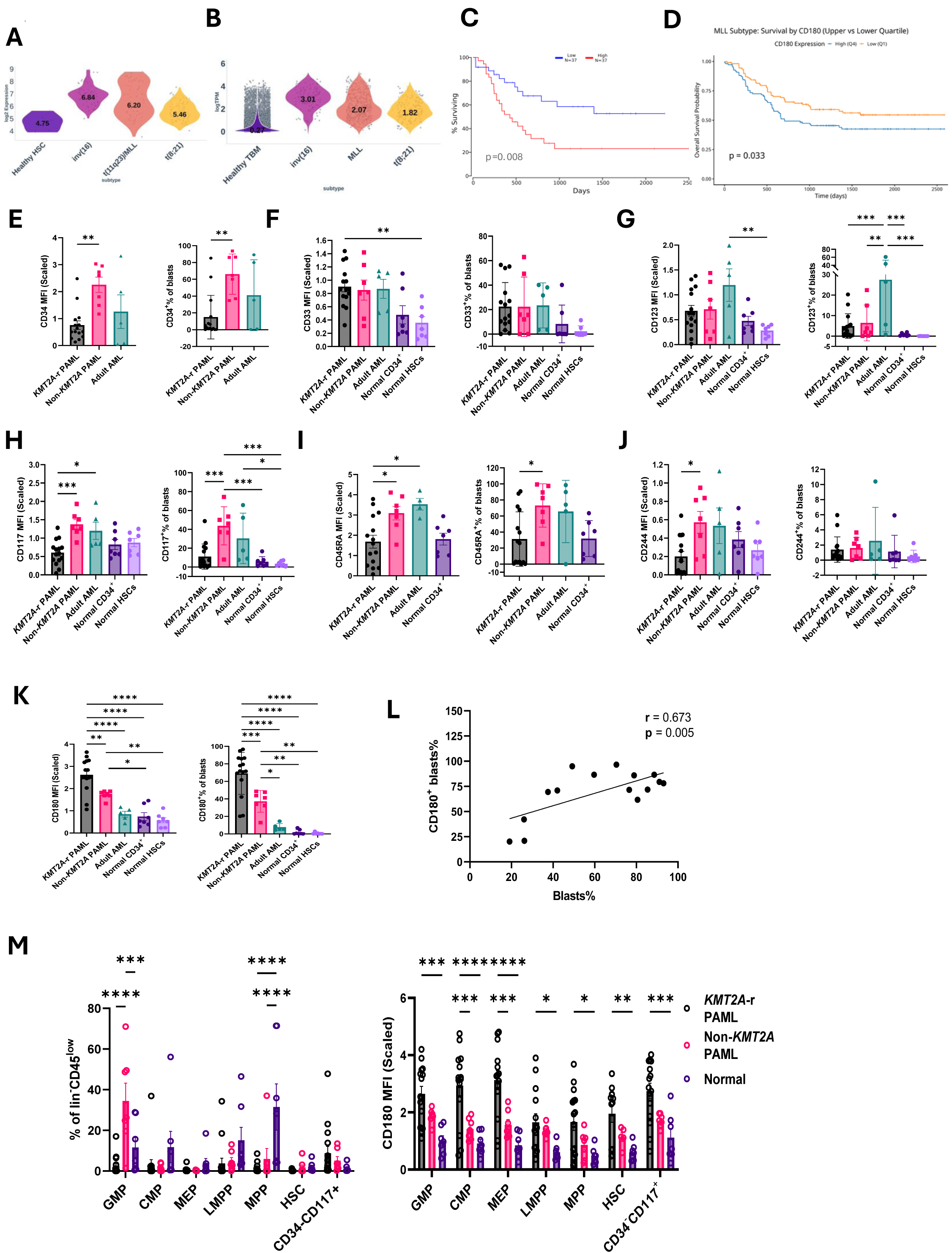
CD180 expression enriched in PAML KMT2A-r. **A-B)** Violin plots of *CD180* gene expression in healthy haematopoietic stem cells (HSC) and total bone marrow (TBM) and AML subtypes with medians shown from **A)** TCGA-LAML and **B)** TARGET and healthy GTEx datasets. **C-D)** Survival curves of the upper and lower quartiles of *CD180* gene expression from **C)** TCGA-LAML dataset, data from http://www.oncolnc.org, and **D)** KMT2A-r patients in TARGET dataset with p value highlighted. **E–K)** Bar plots depicting the mean arcsinh-transformed fluorescence intensity (scaled MFI; left) and the percentage of positive cells (right) for CD34, CD33, CD123, CD117, CD45RA, CD244, and CD180 in blasts from KMT2A-r PAML (n=15), non-KMT2A PAML (n=7), adult AML (n=5), normal CD34^⁺^ cells, and normal haematopoietic stem cells (HSCs) (n=7). One-way ANOVA performed. **L)** Linear regression analysis graph showing the correlation between CD180 and percentage blasts in KMT2A-r PAML diagnostic samples (n=15) with Pearson correlation coefficient (r) and p value indicated. **M)** Bar plots showing (left) the percentage composition of total blasts and (right) CD180 expression levels of the indicated immunophenotypically defined subpopulations (granulocyte-monocyte progenitors (GMP), common myeloid progenitors (CMP), megakaryocyte-erythroid progenitors (MEP), lymphoid-primed multipotent progenitors (LMPP), multipotent progenitors (MPPs), HSC, and CD34-CD117+), in KMT2A-r cases, GMP, MPP, and CD34^⁻^CD117^⁺^ (n=15), CMP, MEP, and LMPP (n=14), and HSC (n=12); in non-KMT2A cases, all subpopulations (n=7) except HSC (n=5); and in normal samples, all subpopulations (n=7). Two-way ANOVA performed.

### CD180 is a marker of leukaemia stemness

To explore *CD180* gene expression and molecular heterogeneity in KMT2A-r PAML, we performed single-cell RNA-sequencing (scRNA-seq) on diagnostic samples from four KMT2A::MLLT3 patients (**Table S1**). BM mononuclear cells (MNCs) were sorted for lineage-negative (lin-; CD3, CD4, CD8, CD19, CD20) CD45^low^ cells to enrich for leukaemic blast cells. These samples all had <2% CD34+ expression, one sample which contained 6.3% CD34+ cells was sorted for CD34-. We created an integrated model from the four samples, comprising a total of 16031 cells, and performed dimensionality reduction and unsupervised clustering followed by UMAP embeddings, resulting in identification of 15 distinct clusters (**Figure 2A**). Feature plots show minimal *CD34* gene expression (0.7% of cells). *CD180 is* expressed in ∼30% cells (4796 cells), with cluster 0 (the largest cluster), 4, 7 and 1 expressing the highest levels of *CD180* (**Figure 2B-D, Figure S2A-B**). To assess stemness, we applied the AML LSC17 gene expression score, that is linked to worse outcomes (26, 27), which showed strong enrichment in *CD180^high^*clusters (0, 1, 4 and 7) and *CD34^high^CD180^low^* cluster 10, compared to *CD180^low^* clusters 5 and 11 (**Figure 2E-F).** Applying the PAML specific LSC6 score (28) and the recently developed PC2-34 score (a score containing 34 genes not including *CD34* gene) for both aAML and PAML (5), also showed that *CD180^high^* clusters have stronger enrichment compared to *CD180^low^* clusters (**Figure S2C-F**). Notably the PC2-34 score which does not contain *CD34* gene (whereas LSC17 and LSC6 do), showed better association with *CD180^high^* clusters. Gene set enrichment analysis using the MSigDB hallmark gene set revealed significant enrichment of the oxidative phosphorylation (OXPHOS) pathway (**Figure 2G**), a process associated with chemoresistance and LSC/LRC persistence in AML (29, 30), and linked to *CD180^high^* clusters at diagnosis (0 and 1). *CD180^high^* clusters 4 and 7 had enrichment of cell cycle related pathways such as E2F and G2M in addition to OXPHOS (**Figure S2G-H**). To assess whether *CD180* is co-expressed with other LSC/LRC-associated genes at the single-cell transcriptomic level, we performed hierarchical clustering using a curated panel of LSC/LRC-associated surface antigens and genes with functional relevance to LSC biology (**Table S2**). *CD180*^high^ clusters (0 and 1) showed distinct co-expression of multiple LSC markers, with cluster 0 and 1 notably expressing *KIT* (CD117), *CD33*, *IL3RA* (CD123) and *CD93* (**Figure 2H**). Correlation analysis within all clusters confirmed strong positive associations between *CD180* and *CD9* and *CD44*, and negative correlations with *CD34* (**Figure 2I**). Directly assessing the stemness functionality of CD180 expressing cells, we performed patient-derived-xenografts (pdx) using primary AML samples (1 PAML, 2 aAML). After 16 weeks, CD180+ cells showed increased frequency and MFI compared to other LSC/LRC markers (**Figure 2J-K**), indicating selective expansion. Xenografts of flow sorted CD180^high^ THP-1 cells exhibited greater engraftment to the bone marrow and brain than CD180^low^ counterparts further confirming *in vivo* leukaemia function of CD180+ cells (**Figure S2I-K**). Validating these data, CD180-knockout (KO) THP-1 cells exhibited significantly less engraftment compared to non-targeting control (NTC) transduced cells (**Figure 2L, 5J-K, Figure S2L-M**). Together, these findings support CD180 as an LSC/LRC-associated antigen linked to stemness-related transcriptional programs.

**Figure 2:**
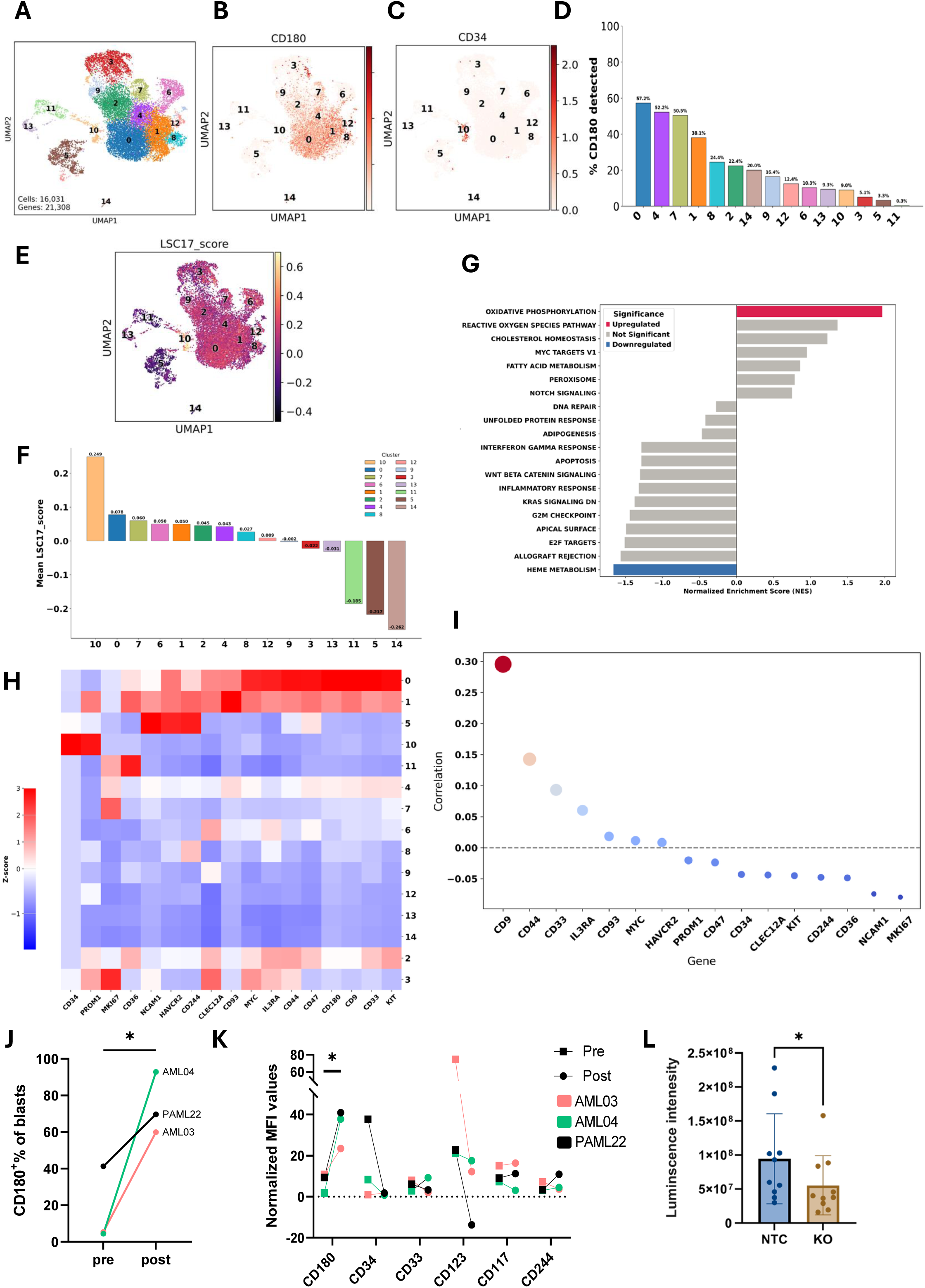
CD180+ marks stem-like cells enriched KMT2A::MLLT3 cells. **A)** UMAP showing 4 diagnosis KMT2A::MLLT3 patients at a Leiden Clustering resolution of 0.6, made up of 16,031 cells and 21,308 genes. **B-C)** Feature plots showing *CD180* **B)** and *CD34* **C)** gene expression in all cell populations. **D)** Bar chart showing percentage of *CD180* expressing cells per cluster. **E)** UMAP showing the LSC17 score across the cell populations. **F)** Box plot quantification of LSC17 score across clusters. **G)** Stacked bar plot showing output of GSEA, where clusters 0 and 1 were compared against all other clusters. Pathways are coloured according to their significance. **H)** Heatmap of relative expression levels of genes related to stemness with the addition of *MKI67* and *CD180*, across each cluster. Expression values were standardized using Z-scores, calculated per gene across all samples. Red indicates higher expression (positive Z-score), blue indicates lower expression (negative Z-score), and white represents average expression (Z-score ≈ 0). Hierarchical clustering was applied to both genes and samples to reveal patterns of co-expression and sample similarity. **I)** Bubble plot illustrating Pearson’s correlation coefficients between CD180 and the genes shown across the entire population of cells. Each bubble represents a pairwise correlation, with colour indicating the strength and direction of the correlation (red for positive, blue for negative). Bubble size corresponds to the statistical significance (p-value), with larger bubbles indicating stronger significance. **J)** Connected line plot showing the percentage of CD180+ blasts in individual patient samples before (pre) and after (post) mouse engraftment. **K)** Connected line plot showing changes in marker expression as geometric MFI normalized to corresponding FMO controls. **L)** Bar graphs depicting quantification of signal intensity at day 17 after transplant with CD180-KO and NTC THP-1 cells. Graphs represent mean ± SD.

CD180’s normal haematopoietic expression profile is not well known. Cord blood analysis showed CD180 mainly on mature B cells, DCs, and monocytes, and near absent haematopoietic stem and progenitor cells (HSPCs) (**Figure 3A**). The expression level of CD180 on KMTA-r PAML blasts was higher in comparison to all healthy populations. scRNA-seq analysis of fetal, paediatric, and adult BM revealed higher *CD180* in FBM/PBM than ABM, largely from monocytes, DCs, and B progenitors (**Figure 3B**). Direct comparison confirmed KMT2A::MLLT3 blasts expressed more CD180 than healthy B cells, monocytes, DCs, and HSPCs (**Figure 3C, S3A-B**). Comparison of cell surface CD180 expression on lineage-positive (lin+) lymphoid cells (CD3, CD4, CD8, CD19, CD20) showed no significant differences across groups (**Figure 3D**). These findings support CD180 as a relatively specific marker for KMT2A-r PAML, despite lymphoid expression, suggesting it may be a direct KMT2A-r target.

**Figure 3:**
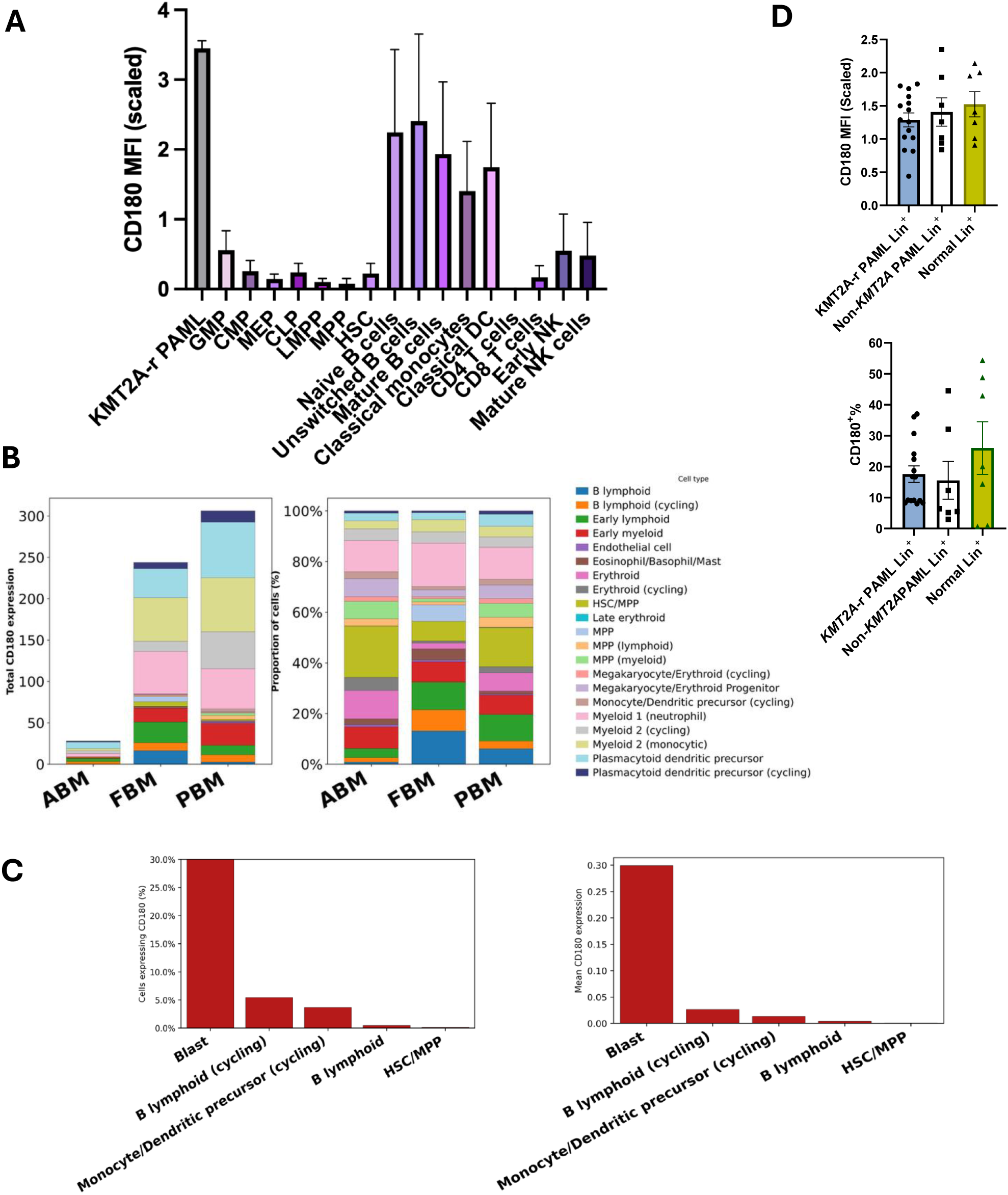
CD180 expression profile across healthy cell ontogeny compared to KMT2A PAML. **A)** Bar plot showing the mean arcsinh-transformed fluorescence intensity (scaled MFI) CD180 expression levels of the indicated immunophenotypically defined subpopulations in cord blood compared to KMT2A-r PAML blasts (n=3). **B)** Bar plots showing the total *CD180* expression in adult bone marrow (ABM), foetal bone marrow (FBM) and paediatric bone marrow (PBM) across the indicated haematopoietic cell type (left), and the proportional composition of each cellular populations in the respective samples (right). Data from publicly available sc-RNA-seq datasets. **C)** Bar plot showing the percentage of cells expressing *CD180* (left) and the mean expression (right), comparing KMT2A-r PAML blasts and selected populations of healthy cells from PBM. **D)** Bar plots depicting the mean arcsinh-transformed fluorescence intensity (scaled MFI; top) and the percentage of CD180+ cells (right) in lineage-positive (lin+) cells, defined as CD3^⁺^, CD4^⁺^, CD8^⁺^, CD19^⁺^, or CD20^⁺^, comparing KMT2A-r PAML (n=15), non-KMT2A PAML (n=7), and normal samples (n=7). Cell populations include haematopoietic stem cells (HSC), common myeloid progenitors (CMP), granulocyte-monocyte progenitors (GMP), common lymphoid progenitors (CLP), megakaryocyte-erythroid progenitors (MEP), multipotent progenitors (MPPs), lymphoid-primed multipotent progenitors (LMPP), unswitched B cells, naive B cells, mature B cells, classical monocytes, classical dendritic cells (Classical DC), CD4 T cells, CD8 T cells, early natural killer cells (Early NK cells), and mature natural killer cells (Mature NK cells).

### CD180 is a direct target of KMT2A::MLLT3

KMT2A::MLLT3 interacts with various transcriptional factors and chromatin-modifying complexes to activate genes critical for maintaining stem cell properties and promoting the development of leukaemia (31, 32). Firstly, FISH analysis confirmed the leukaemic nature of CD180 expressing cells, with detection of the KMT2A::MLLT3 translocation in a significantly higher proportion of flow sorted CD180+ patient blasts compared to CD180- cells (**Figure 4A-B**) (an average of 63.91% versus 26.45%, respectively), and validated using THP1 and U937 cell lines as positive and negative controls (**Figure S4A**). To validate leukaemic transcriptional activity in CD180+ blast clusters, we defined a KMT2A::MLLT3 target gene signature, starting with a core set of KMT2A::MLLT3-responsive genes (33) and expanded by integrating essential KMT2A-r targets (34–36) (**Table S3**). Using Scanpy’s in-built function (Score_genes), we quantified per-cell transcriptional activity at diagnosis, revealing strong enrichment of the KMT2A::MLLT3 gene score in *CD180^high^* clusters (0, 4, 7 and 1) versus *CD180^low^* clusters (13, 5, and 11). Hierarchical clustering confirmed this pattern, with *CD180^high^* clusters showing consistently elevated KMT2A::MLLT3 gene score, while *CD180^low^* clusters (13, 5, and 11) showed minimal activity. Notably, cluster 10 was the only cluster that expressed CD34 and exhibited a *CD180^low^*LSC17^high^KMT2A^low^ profile, suggesting a distinct transcriptional profile for CD34+ cells in these patients (**Figure 4C-E, Figure S4B**). These findings support CD180 as a marker of the leukaemic clone in KMT2A::MLLT3 PAML and possibly a direct target gene of the complex. To address this possibility, we conducted analysis of KMT2A::MLLT3+ THP-1 AML cell line CHIP-seq data (37, 38). We identified KMT2A binding at two intragenic regions of the *CD180* locus, one within intron 1 and another at the 3′ untranslated region (UTR). These regions also showed co-occupancy by MLLT3 (AF9), consistent with binding by the KMT2A::MLLT3 fusion complex (**Figure 4F**). Chromatin at these sites exhibited features of active enhancers, including enrichment for H3K79me3, H3K27ac, H3K4me1, and absence of the repressive mark H3K27me3 (track not included). BRD4 was also bound at these regions, suggesting transcriptional coactivation. To further investigate the regulatory potential of these intragenic elements, we performed ATAC-seq and RNA-seq on THP1 cells sorted into CD180^high^ and CD180^low^ populations. These analyses revealed that the intragenic regions were associated with open chromatin in CD180^high^ cells and coincided with increased *CD180* transcript levels, supporting their role as active regulatory elements (**Figure 4F)**. Previous studies identified CD180 as a pharmacodynamic biomarker of BET inhibitors (targeting BRD4) (39, 40), with promising pre-clinical data although clinical trial data suggest mixed efficacy of these agents in AML (41). Menin inhibitors, which block the KMT2A complex from activating downstream target genes, have shown promising results in clinical trials (42). To validate KMT2A::MLLT3 complex-mediated regulation of CD180, we treated AML cells with Menin (Revumenib) and BET inhibitors (AZD5153) and observed downregulation of CD180 expression in KMT2A::MLLT3 human cell lines (**Figure 4G-J)**. Together, these findings confirm *CD180* as a direct transcriptional target of KMT2A::MLLT3 and suggest the presence of two intragenic fusion-bound enhancers that regulate its expression.

**Figure 4:**
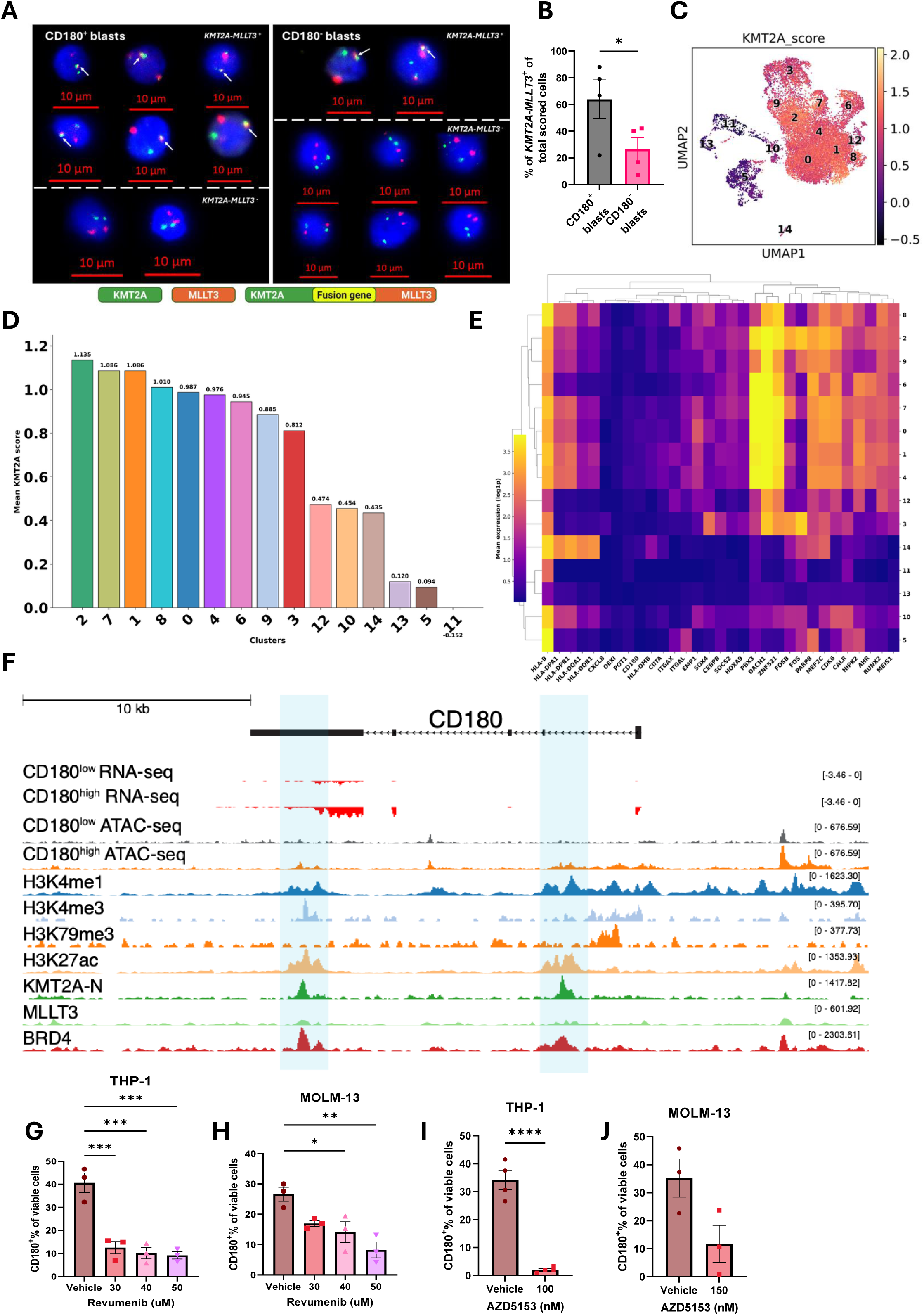
CD180 marks KMT2A::MLLT3 leukemic blasts and shows direct regulation by the fusion complex. **A)** Representative Fluorescence in situ hybridization (FISH) images of nuclei from sorted CD180+ (left) and CD180- (right) PAML blasts showing the KMT2A::MLLT3 translocation signal (yellow, indicated by white arrows), along with distinct KMT2A (green) and MLLT3 (orange) signals. Nuclei are counterstained with DAPI. Images were acquired at 63× magnification, and scale bars are shown. **B)** Bar plot depicting the percentage of *KMT2A*::*MLLT3+* fusion signals scored in sorted CD180+ and CD180- blasts (n = 4). **C)** UMAP plot of the defined KMT2A score in the 4 diagnosis KMT2A::MLLT3 clustered populations. **D)** Bar plot of the mean quantification of the KMT2A score across the clusters. **E)** Heatmap showing the mean expression of all the KMT2A genes in the KMT2A score, and *CD180,* across the clusters. Log transformed gene expression values shown (high is yellow, low is purple). **F)** ChIP-seq tracks for KMT2A, MLLT3, H3K4me1, H3K4me3, H3K79me3, H3k27ac and BRD4 together with the ATAC-seq and RNA-seq tracks from sorted THP1 CD180^high^ vs. CD180^low^ populations at the *CD180* locus. KMT2A occupancy seen at intron 1 and 3′ UTR, with co-occupancy by MLLT3, with active enhancer marks (H3K79me3, H3K27ac, H3K4me1) and BRD4 binding, open chromatin and increased CD180 transcript levels in CD180^high^ cells. **G-H)** Bar plots showing the percentage of CD180+ viable cells following treatment with different doses of the menin inhibitor Revumenib in THP-1 **G)** and MOLM-13 **H). I-J)** Bar plots showing the percentage of CD180+ viable cells following treatment with bromodomain and extra-terminal domain inhibitor (BETi) AZD5153 in THP-1 **G)** and MOLM-13 **H).**

### CD180 marks quiescent cells

*Ki-67* expression showed a distinct negative correlation with *CD180,* and specifically in *CD180*^high^ clusters (0 and 1) (**Figure 2H-I**). This suggests a quiescent phenotype which is linked to LSC/LRC features and chemoresistance. UMAP and cell cycle phase analysis revealed that the majority of *CD180^high^* cells are *ki-67^low^* (0 and 1) with G1 cell cycle features. However, there are *CD180^high^* clusters that are *ki-67^high^*clusters (4 and 7) with S and G2M features (**Figure 5A-D, Figure S5A**). Flow sorted CD180^high^ THP-1 and Molm-13 cells exhibited slower proliferation, and more cells in G0 measured by flow cytometry with Ki-67 and DAPI staining, than CD180^low^ counterparts in culture (**Figure 5E-H**). Primary AML patient samples sorted by CD180 expression revealed a higher proportion of cells in G0 phase within the CD180^high^ population (**Figure 5I**). Validating these data, CD180-KO THP-1 cells exhibited significantly less G0 quiescent cells compared to NTC transduced cells, without any effect on cell viability (**Figure 5J-L**). Furthermore, overexpression (OE) of CD180 in THP-1 cells led to significantly more G0 quiescent cells compared to empty vector control (EV) transduced cells (**Figure 5M-O, Figure S5B**). Together, these findings support molecular characterisation of a CD180-associated quiescent phenotype.

**Figure 5:**
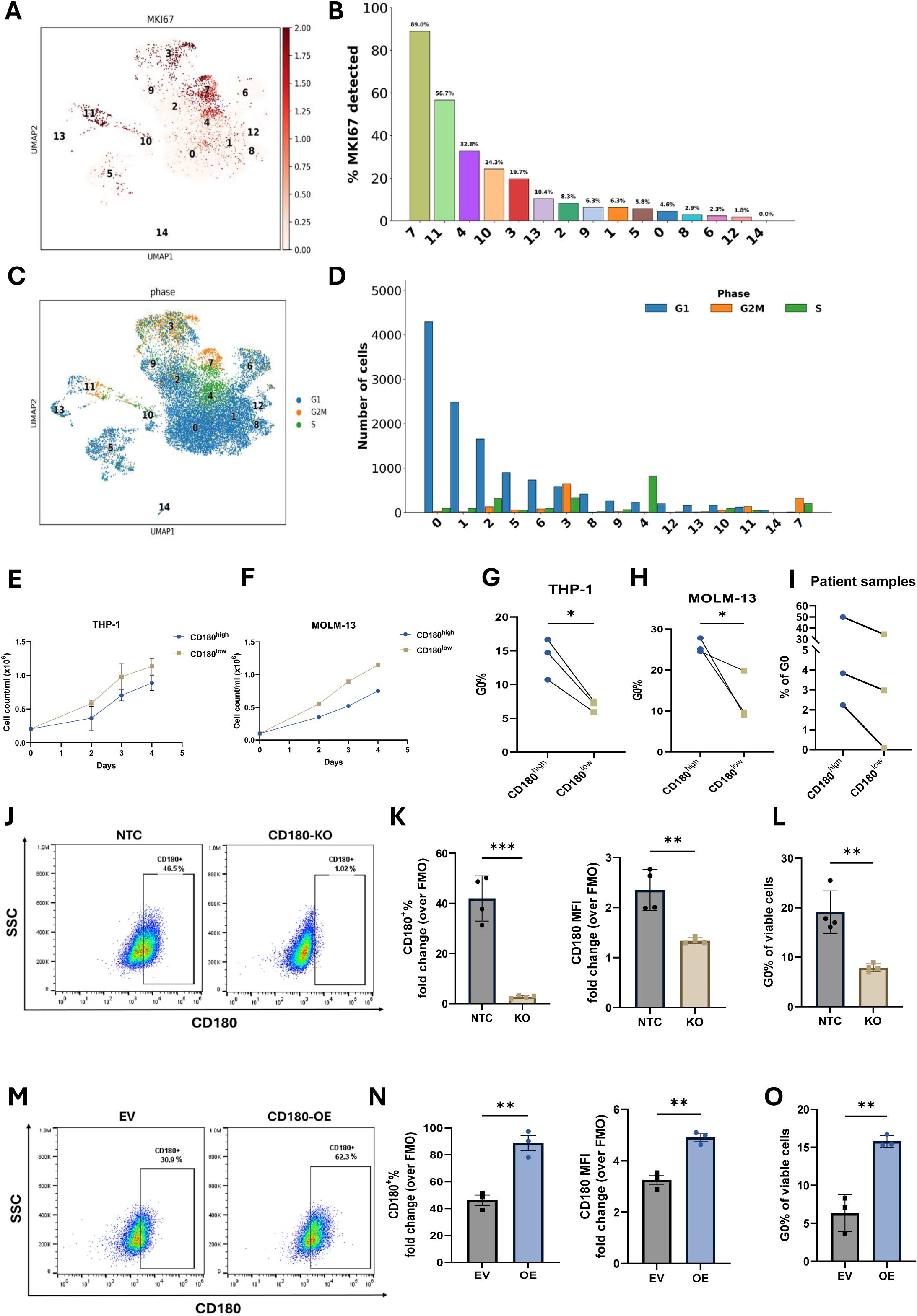
CD180 expression associates with a quiescent phenotype and cell cycle state in KMT2A::MLLT3 PAML. **A)** UMAP plot of *MKI67* expression in the 4 diagnosis KMT2A::MLLT3 clustered populations. **B)** Bar plot showing the percentage of *MKI67* expressing cells across clusters. **C-D)** UMAP plot of cell cycle phases **C)** and a grouped bar plot showing the number of cells in each cluster in each cell cycle phase **D)** in the 4 diagnosis KMT2A::MLLT3 clustered populations. **E-F)** Cell growth curves over time of CD180^high^ and CD180^low^ sorted THP-1 (n = 3) **E)** and MOLM-13 (n = 2) **F). G-H)** Bar plots depicting the percentage of G0 quiescent cells in CD180^high^ and CD180^low^ sorted THP-1 **G)** and MOLM-13 **H)** cells. **I)** Connected line plot showing the percentage of G0 cells in CD180^high^ and CD180^low^ diagnostic blasts from patient samples; PAML07, PAML14, and AML01. Lines connect paired measurements. **J and M)** Representative flow cytometry dot plots showing CD180 surface expression of genetically edited THP-1 cells non-targeting control (NTC, left) and CD180 knockout (CD180-KO, right) **J)** and in empty vector (EV, left) versus CD180 overexpression (CD180-OE, right) **M).** Gates are based on the FMO controls. **K and N)** Bar plots showing CD180 expression in CD180-KO versus NTC cells (n = 4) **K)** and CD180-OE versus EV control cells (n = 3) **N).** Expression is quantified as the percentage of CD180^⁺^ live cells (left) and CD180 MFI (right), with both measures normalised to the respective FMO controls. **L and O)** Bar plots showing the percentage of viable cells in G0 phase in CD180-KO versus NTC cells (n = 4) **L)** and CD180-OE versus EV control cells (n = 3) **O).**

### CD180 associates with chemoresistance and relapse

We examined CD180’s role in chemoresistance. CD180 was upregulated in THP-1 and MOLM-13 cells after AraC and Mitoxantrone treatment. Chemoresistant lines (AraC- and Mitoxantrone-resistant) showed markedly higher CD180 than isogenic parental cells. Sorted CD180^high^ cells were significantly more drug-resistant than CD180^low^ cells, with higher EC50 and viability (**Figure 6A–I and Figure S4C-D, S6A-B**. CD180 KO cells exhibited significantly reduced chemoresistance relative to controls (**Figure 6J**), while CD180 OE cells showed elevated resistance (**Figure 6K**), further implicating CD180 in modulating drug responses. To address the chemoresistance in the context of persistence of CD180 expressing cells in patients undergoing treatment, we assessed CD180 profiles in KMT2A::MLLT3 patients in the Myechild 01 clinical trial from matched diagnosis (Dx), post-course 1 (PC1), and relapse samples (Rx). The total percentage of blast cells decreased at PC1 and returned at Rx, and consistently the percentage of CD180+ blasts mirrors this pattern (**Figure 6L-M**). Analysis of the MyeChild 01 clinical data for PAML patients (including all samples in our cohort) revealed no correlation between CD180 MFI or blast percentage with age (**Figure 6P, Figures S6J-L**). CD180 expression correlated negatively with hemoglobin and platelet counts and positively with white blood cell count. No significant differences were observed between relapsed and non-relapsed diagnostic samples (**Figure 6Q-R; Figures S6M-P**). Focused analysis on LSC/LRC potential populations driving Rx in these patients showed that while no prominent CD34+CD38- population was evident at Rx (**Figure 6N**), a return of a CD34-CD117+CD180+ population was detected **(Figure 6O**, **Figure S6E-I)**. However the limiting number of matched Rx cells/samples in our flow analysis preclude us from concluding whether a more refined LSC/LRC marked population was prominent at Rx. To investigate phenotypic changes in KMT2A::MLLT3 patient blasts over disease progression, we applied the FlowSOM algorithm, based on the flow cytometry data, for unsupervised clustering of matched samples at Dx, PC1, and Rx. FlowSOM identified 21 distinct clusters, revealing increasing phenotypic heterogeneity/shifting from Dx to Rx. Notably, cluster 12, which was sparsely represented at Dx, expanded at PC1 and further at Rx. This cluster showed significantly higher CD180 expression compared to other LSC/LRC-associated antigens and was consistently negative for CD34 (**Figure 6S-U)**. These data support CD180 as a marker of treatment resistant cells that associates with relapse.

**Figure 6:**
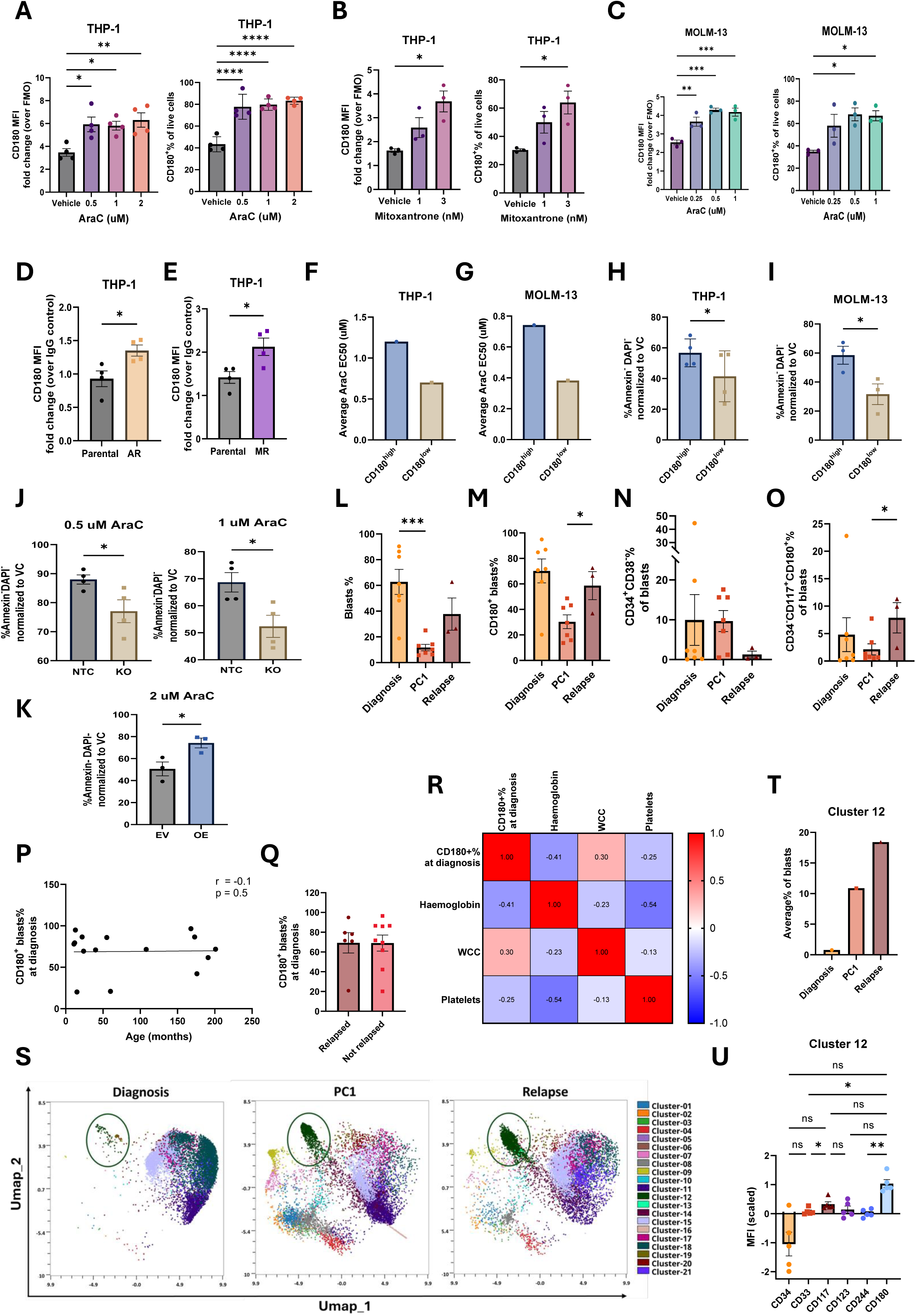
CD180 expression associates with chemoresistance and persistence in KMT2A::MLLT3 PAML. **A-C)** Bar plots depicting CD180 MFI normalized to the corresponding FMO control in THP-1 cells treated with the indicated doses of cytarabine (AraC) **A),** mitoxantrone **B),** and AraC in MOLM-13 cells **C)** (n = 4). **D-E)** Bar plots depicting CD180 MFI expression in parental THP-1 cells compared to isogenic AraC-resistant (AR) and mitoxantrone-resistant (MR) THP-1 cells (n = 4). **F-G)** Bar plots showing average EC50 values for AraC in CD180^high^ and CD180^low^ sorted THP-1 (n = 4) and MOLM-13 (n = 3), respectively. **H-I)** Bar plots showing the percentage of viable cells (Annexin^−^DAPI^−^) in CD180^high^ and CD180^low^ sorted THP-1 cells treated with 1 uM AraC **H),** and MOLM-13 cells treated with 0.5 uM AraC **I). J)** Bar plots showing the percentage of viable cells (Annexin^−^DAPI^−^) normalized to vehicle control (VC) in CD180 knock-out (CD180-KO) THP-1 cells compared to non-targeting control (NTC) cells following treatment with 0.5 uM (left) or 1 uM (right) AraC for 48 hours (n = 4). **K)** Bar plot showing the percentage of viable cells (Annexin-DAPI-) in CD180 overexpressing (CD180-OE) THP-1 cells compared to the empty vector (EV) control following treatment with 2uM AraC. **L)** Bar plot depicting changes in blast cell percentages across disease stages (diagnosis (n=7), post–course 1 (PC1, n=7), and relapse (n=3)) in KMT2A-r PAML samples. **M-O)** Bar plots depicting changes in **M)** the percentage of CD180^⁺^ blasts, **N)** the percentage of CD34-CD117+CD180+ blasts, and **O)** the percentage of CD34+CD38- blasts across disease stages in KMT2A-r PAML. **P)** Correlation between the percentage of CD180^⁺^ blasts at diagnosis in KMT2A-r PAML (n = 15) and patient age (months). Scatter plot shows the linear regression line with Pearson correlation coefficient (r) and p value indicated. **Q)** Bar plot showing the percentage of CD180+ blasts at diagnosis in KMT2A-r PAML patients who relapsed (n = 6) and in patients who did not relapse by the time of analysis (n = 9). **R)** Correlation matrix of the percentage of CD180+ blasts at diagnosis in KMTA-r PAML with patient haematological parameters including haemoglobin, white cell count (WCC), and platelets (n = 10). Pearson correlation coefficients indicated, with red and blue shading representing strong positive and negative correlations, respectively. **S)** Flow Self-Organizing Map (FlowSOM) clustered UMAPs depicting the different clusters assigned based on the phenotypic profiles of *KMT2A*-r PAML blasts at the different disease stages. Cluster 12 is highlighted with dark green circles. Two matched *KMT2A*-r PAML patient samples were used in this analysis. **T)** Bar plot depicting the average percentage of cluster 12 of the total blasts in *KMT2A*-r PAML different stages. **U)** Bar plot showing CD34, CD33, CD117, CD123, CD244, and CD180 expression in cluster 12 across all samples used (2 matched samples, n = 5 due to absence of cluster 12 in one of the diagnostic samples).

### CD180 clusters exhibit phenotypic shifts during disease progression

To characterise *CD180* expressing cells during progression, scRNA-seq was performed on matched Dx (n=4), PC1 (n=4), and Rx (n=2) samples (lin- CD45^low^ cells, 57,874 cells, 23,629 genes). Integration revealed 18 clusters (**Figure 7A-B**). UMAP showed dynamic *CD180* expression patterns. At relapse, clusters 0, 8, 11, and 14 contained most *CD180^high^* cells (**Figure 7C-D, Figure S7A**). Cluster 0 reappeared at Rx after absence at PC1, cluster 14 declined then returned, while clusters 8 and 11 persisted and expanded, indicating phenotypic shifts and survival of chemoresistant clones (**Figure 7E**). Dx-PC1-Rx *CD180^high^* clusters (e.g. 0, 8, 11, and 14) had high KMT2A-gene target and LSC17 scores compared to *CD180^low^* clusters (e.g. 1, 2, 17) (**Figure 7F-I**). The PC2-34 and PAML LSC6 score, again, associated with *CD180^high^* expressing clusters (**Figure 7J-K, Figure S7B-C**. Notably, Dx-PC1-Rx *CD180^high^* clusters include both *ki-67^low^* (clusters 0 and 8) and *ki-67^high^* (clusters 11 and 14) populations (**Figure 7L, Figure S7D**). We assessed OXPHOS activity across Dx-PC1-Rx clusters. It remained a feature of *CD180^high^* clusters but was less pronounced at relapse (**Figure 7M, Figure S7E**). GSEA revealed relapse clusters shifted toward stemness, therapy resistance, immune evasion, metabolic flexibility, and inflammatory pathways (WNT, Notch, Hedgehog, TNF, TGFβ) (**Figure 7N**).

**Figure 7:**
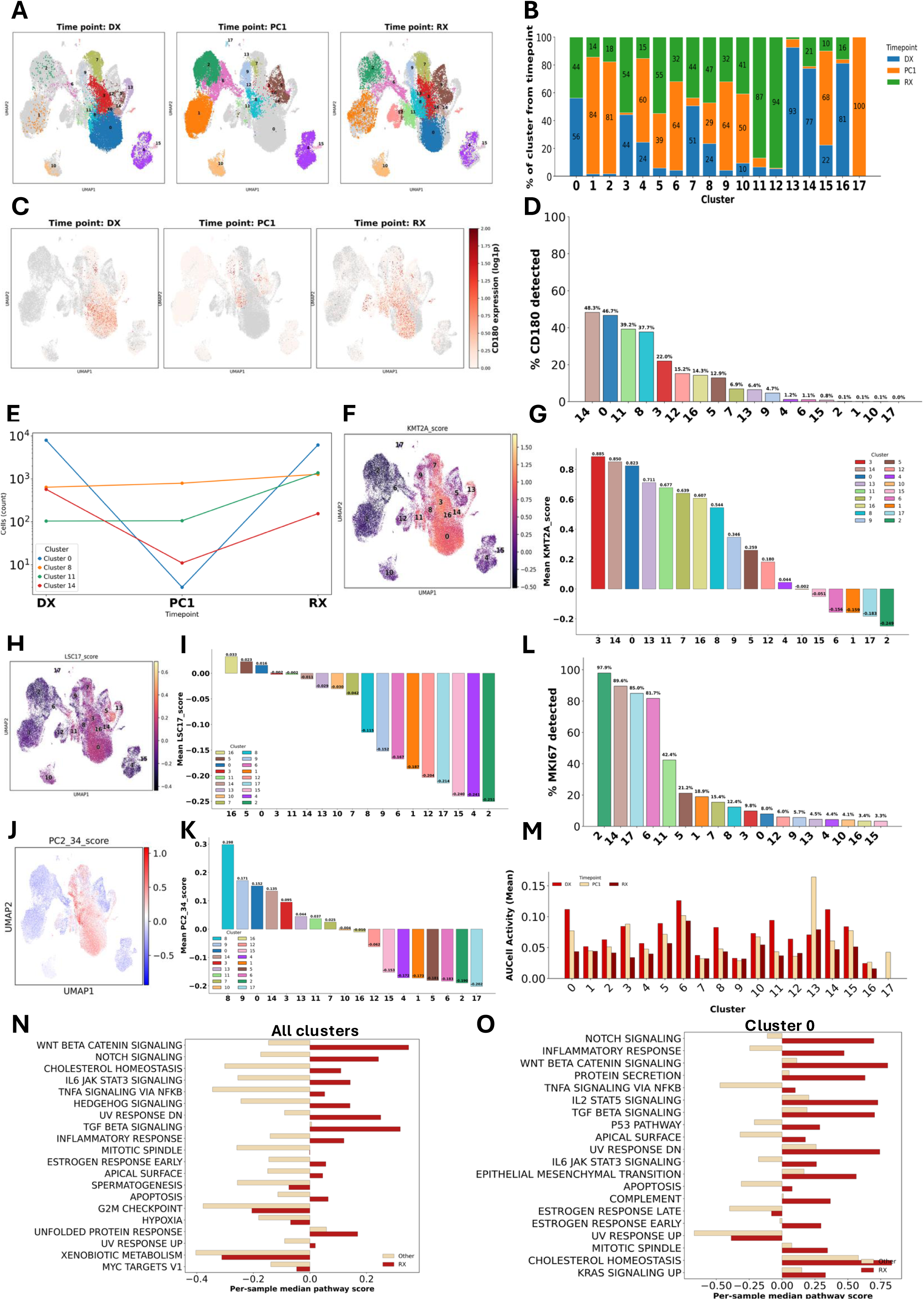
Dynamic evolution of CD180+ subpopulations across disease progression. **A)** An integrated UMAP of matched diagnosis (n=4), PC1 (n=4) and relapse (n=2) KMT2A::MLLT3 patient samples with cells coloured and separated by disease stage. **B)** A stacked bar plot showing the percentage makeup of each cluster by disease stage with each stage assigned a different colour. Percentages labelled on bars. **C)** Integrated UMAP feature plot of *CD180* gene expression in all cell populations. **D)** Bar graph showing the percentage of *CD180* expressing cells per cluster. **E)** Line graph tracking the number of cells at the different stages for cluster 0, 8, 11 and 14. **F-K)** UMAPs **F and H and J)** and bar graphs **G and I and K)** of KMT2A and LSC17 and PC2_34 mean scores respectively for each cluster coloured according to the integrated UMAP. **L)** Bar plot showing the percentage of *MKI67* expressing cells across clusters. **M)** Grouped bar plot showing the mean AUCell activity score for oxidative phosphorylation activity across each of the clusters at the different disease stages. **N-O)** Bar plot showing the per sample median for the combined pathway activity score for relapse vs other disease stages across the whole cell population **N)** and relapse cluster 0 vs other disease stages cluster 0 **O).**

Comparing *CD180^high^* cluster 0 at relapse vs diagnosis revealed a more stem-like, chemoresistant, and inflammatory phenotype at relapse (**Figure 7O).** Persisting *CD180^high^* clusters 8, 11, and 14 expanded at relapse, showing enhanced stemness and metabolic features (**Figure S7F-H**). These findings show that *CD180^high^* subpopulations shift dynamically during progression, acquiring stem-like, chemoresistant, and metabolically adaptive traits, implicating them in relapse and as therapeutic targets. Clusters absent at PC1 but reappearing at Rx, and others persisting and expanding, indicate clonal evolution and phenotypic plasticity. The longitudinal analysis identifies *CD180^high^* cells as key relapse mediators linked to stemness, therapy resistance, and metabolic adaptation.

## Discussion

This study identifies CD180 as a novel and functionally relevant marker associated with chemoresistant cells and relapse in KMT2A-r PAML. Our findings demonstrate that CD180 is not only enriched on leukaemic blasts but also marks distinct LSC/LRC-like populations that persist through treatment and relapse. The data suggests KMT2A-r PAML can be classified as CD34- PAML. This observation parallels recent findings in adult AML, where Pei et al. identified monocytic LSCs that are also CD34-negative (43), and Zeisig et al. showed that humanised KMT2A-r AML model reside in CD34-/lo compartments (44), reinforcing the concept that relapse-driving LSC-like cells can reside outside conventional CD34+ compartments (11). Our study advances this paradigm by identifying CD180 as a mechanistically distinct biomarker in paediatric KMT2A-rearranged AML. Whilst we cannot pinpoint exact immunophenotypic “LSC” subpopulations, collectively, these findings reveal that KMT2A-r PAML displays a distinct immunophenotypic surface antigen profile, characterised by reduced expression of canonical LSC and MRD markers typically used in aAML, while CD180 emerges as a distinguishing and functionally relevant feature. Our analysis revealed that *CD180^high^* clusters were enriched for stemness-associated transcriptional programs, including aAML LSC17 and PAML LSC6 and PC2-34 scores, OXPHOS, and cell cycle regulation. Although the LSC17 and LSC6 scores include CD34, we found that CD180^high^ subpopulations preferentially co-express other LSC-associated genes and are enriched for chemoresistant and quiescent pathways, indicating that stemness signatures are maintained independently of CD34 expression. This highlights the need for refined MRD strategies in PAML, particularly in CD34- subtypes. Our data show that CD180 is broadly upregulated across CD34- and CD34+ (when present) compartments, suggesting it may mark multiple functionally distinct leukaemic subpopulations and overcome limitations of CD34-based detection. Interestingly, one patient with blasts expressing >70% CD180 and <0.5% CD34 and <1% CD33 had CNS relapse. While this observation is intriguing, it suggests a potential association between high CD180 expression and CNS involvement that warrants further investigation.

Mechanistically, we show that CD180 is a direct transcriptional target of the KMT2A::MLLT3 fusion complex, with intragenic enhancer regions bound by KMT2A and MLLT3, and marked by active chromatin features. This is further supported by recent work which demonstrated that KMT2A fusion proteins drive patient-specific enhancer activation in leukaemia (45). The findings implicate KMT2A fusion complexes in the regulation of genes such as MEIS1 and RUNX2 via novel enhancers, and by extension, support our observation of CD180 enhancer regulation by KMT2A::MLLT3. Further work is needed to fully understand how the complex is regulating the epigenetic and transcriptional status of the CD180 locus.

Functionally, *CD180^high^* cells exhibited quiescence and chemoresistance, hallmarks of LSC/LRC, and were enriched in the G0 phase of the cell cycle. CD180 KO did not induce cell death, suggesting that CD180 is dispensable for basal AML cell viability or that alternative survival pathways compensate for its loss. Both KO and OE experiments confirmed that CD180 modulates growth, cell cycle dynamics and chemoresponse, further suggesting CD180 role in maintaining dormancy and survival under therapeutic pressure. These findings were corroborated in pdx and xenograft experiments, where CD180+ populations successfully engrafted, and in our transcriptomic analysis, which showed that *CD180^high^* subpopulations persisted post-treatment and re-emerged at relapse in matched patient samples.

Our longitudinal single-cell analysis revealed that *CD180^high^*subpopulations are not static but dynamically evolve during therapy and relapse. The cluster re-emergence and expansion patterns indicate that CD180 expression marks subclones capable of both clonal evolution and phenotypic plasticity under therapeutic pressure. Mechanistically, these *CD180^high^*clusters exhibit enrichment for stemness-associated pathways, metabolic flexibility, and immune evasion signatures, consistent with a chemoresistant phenotype and mirror features described in therapy-resistant AML subclones in recent single-cell studies (46, 47). The re-emergence and expansion of these populations at relapse suggest survival of therapy-resistant clones that adapt through stem-like programmes and inflammatory signaling. Current state-of-the-art research, predominantly from aAML studies, emphasises that relapse in AML is often driven by rare, therapy-tolerant progenitor-like or quiescent stem-like cells that survive initial chemotherapy and expand under selective pressure. Our findings align with this paradigm, highlighting *CD180^high^*clusters as potential reservoirs of resistance. The enrichment of stemness and immune evasion signatures is consistent with reports linking these traits to MRD and poor outcomes. Furthermore, metabolic adaptability and inflammatory signalling have emerged as hallmarks of aggressive AML phenotypes, reinforcing the clinical relevance of these observations (48, 49). These data position *CD180^high^* clusters within the broader framework of relapse biology and underscore CD180’s potential as a therapeutic target.

Recently a study identified three transcriptionally distinct leukaemia stem and progenitor cell (LSPC) states within AML samples; quiescent, primed, and cycling (5). Quiescent LSPCs, strongly linked to stemness and poor outcomes, expand at relapse and outperformed traditional immunophenotypes (CD34+CD38-) in predicting resistance, while cycling LSPCs also associate with adverse survival. In PAML, relapse is marked by a striking expansion of total LSPCs, particularly the quiescent subset reinforcing their role in therapy resistance. These findings establish quiescent and cycling LSPC signatures as critical determinants of AML hierarchy, prognosis, and relapse biology. Importantly, the study revealed that hierarchy-based gene expression scores predict treatment benefit in adults but not in paediatric AML: adult-derived signatures LinClass-7 and LSC17, both containing CD34, showed limited utility in children, whereas the newly developed 34-gene score (PC2-34), which excludes CD34, accurately captured the Primitive-Mature axis and predicted drug response across both aAML and PAML. This suggested that PAML may require a different biomarker or scoring system to guide therapy selection. Although both our *in vitro* cellular and single-cell transcriptional data showed that CD180^high^ cells mainly represent dormant (G0, *ki-67^low^* ) populations, a few CD180^high^ clusters also exhibited *ki-67^high^* expression, while all *CD180^high^*clusters retained enriched PC2-34 scores. This suggests that CD180 marks LSC/LRC populations spanning both quiescent and cycling states, and targeting CD180 could eliminate both populations, offering a broad therapeutic effect. Indeed, a gene score based on the CD180 transcriptional signature may provide an additional tool for therapy stratification and prediction.

Collectively, these findings identify CD180^high^ cells as key mediators of relapse and position CD180 as a rational target for eradicating leukaemic stem-like reservoirs. Its selective expression on leukaemic cells, coupled with absence from normal HSCs, offers a promising therapeutic window. CD180 also emerges as a clinically actionable biomarker for MRD monitoring, relapse prediction, and therapeutic intervention in KMT2A-r PAML.

## Ethics statement

All animal procedures were conducted in accordance with institutional guidelines and complied with the ARRIVE guidelines for reporting animal research. Experiments were carried out in accordance with the Animals Scientific Procedures Act of 1986 and following the University of Glasgow Animal Welfare and Ethical Review Board under Keeshan’s Home Office Licence (Procedure Project License (PPL) no. PP4496278). The work with patient samples is ethical approved by West of Scotland Research Ethics Service approval 20/WS/0066 and 25/WS/0056. Samples have been approved for release locally by the haematological cell research bank, approved by the VIVO biobank, and approved by the Myechild 01 trial management team.

## Data availability statement

All data associated with this study are present in the paper or the Supplementary Materials. All newly generated datasets supporting this study have been deposited in the Gene Expression Omnibus (GEO) under accession numbers GSE311612, and GSE313999, GSE314000, GSE314001 Series records.

Computational code available at https://zenodo.org/records/17910967?token=eyJhbGciOiJIUzUxMiJ9.eyJpZCI6ImQ4NDg2ZmUxLTQ3ZmEtNDQ5Zi1hMWQyLWEzNjJiZjNlY2JiOCIsImRhdGEiOnt9LCJyYW5kb20iOiJmYTg4NDVhYzA4YzA1N2NkZWFmMDk0YjQ0NTI0ZTQ4NCJ9.72M4LEPo1u7MtJuMUURDEUn28gbp-bz9I3VmAqi0LXv4DKfRi-7rs7QmArXgJCHqYTZMyko-zJ2_zigbXIx-Vw

## Acknowledgments

We acknowledge the facilities, and scientific and technical assistance from flow cytometry staff at Wolfson Wohl Cancer Research Centre and the Biological Services Unit and technical staff at Cancer Research UK Scotland Institute. We are especially grateful for the technical assistance from Amelia Bywater and all members of the Keeshan group. We are grateful to all the investigators in the Myechild 01 clinincal trial and all the patients who consented their samples for research.

## Author Contributions

M.M.E., A.W., F.F.M., E.K., M.T.S., A.L.S., E.R., A.C., A.G., K.M.R., M.A., performed experiments. M.M.E., A.W., and A.L.S., drafted the figures. M.M.E., A.W., F.F.M., E.K., A.L.S., E.R., A.C., T.S., M.A., analysed the data. J.C., K.D., provided expert technical advice and support. G.V.H., T.A.M., B.G., P.V., C.J.H., D.V., provided expert advice and essential resources. L.W., B.G., P.V., and C.J.H provided essential clinical information and resources. K.K. drafted the manuscript with contributions from M.M.E., A.W., and all authors edited and approved the final version. K.K. designed and supervised the project.

