## Supplementary figures 1-7 for "CD180 identifies chemoresistant stem-like blasts and reveals a KMT2A-driven vulnerability in acute myeloid leukaemia"

### Supplementary Figure 1

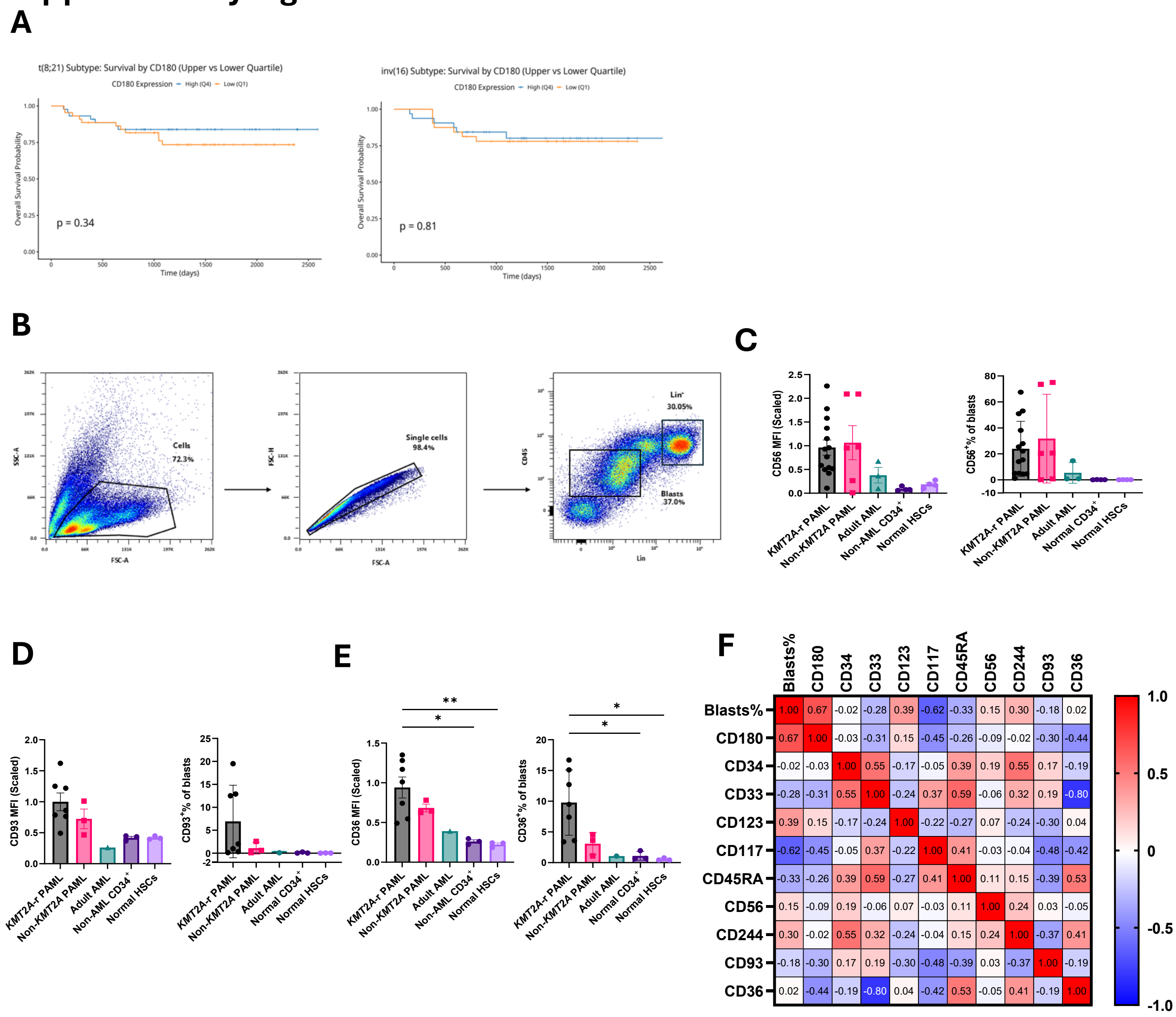

### Supplementary Figure 2

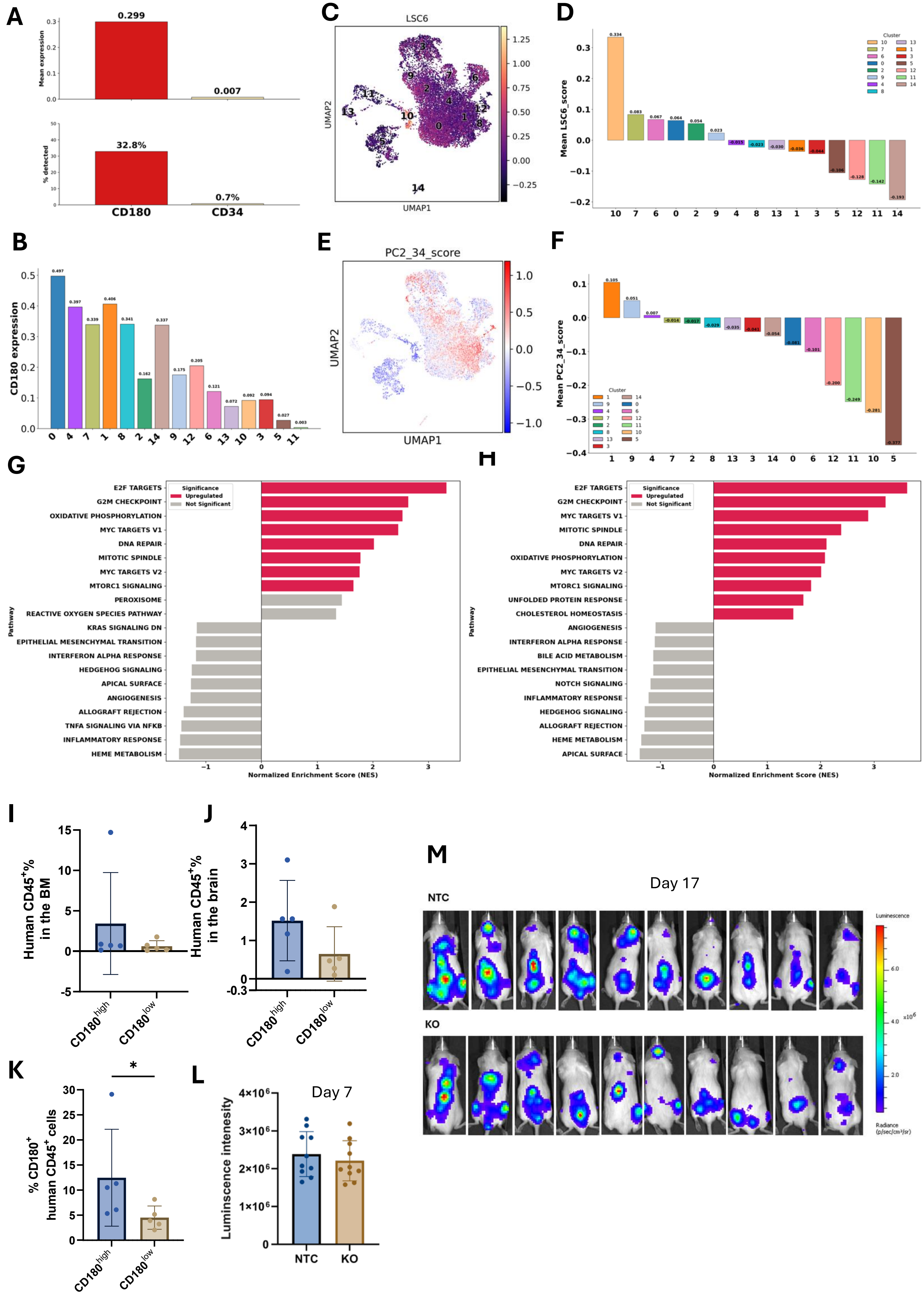

### Supplementary Figure 3

**A**

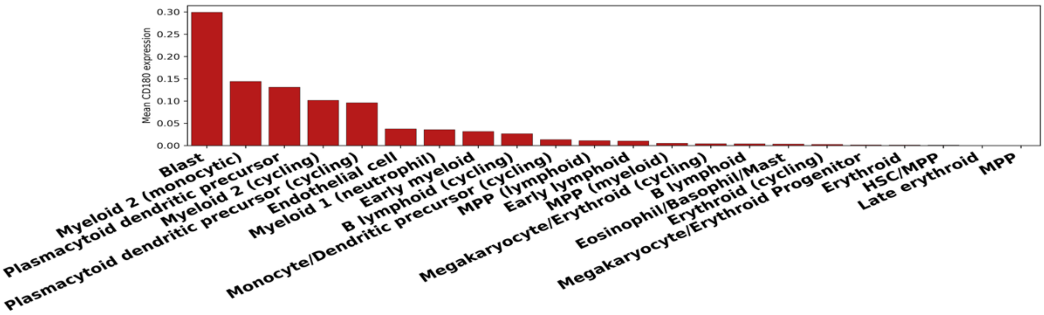


**B**

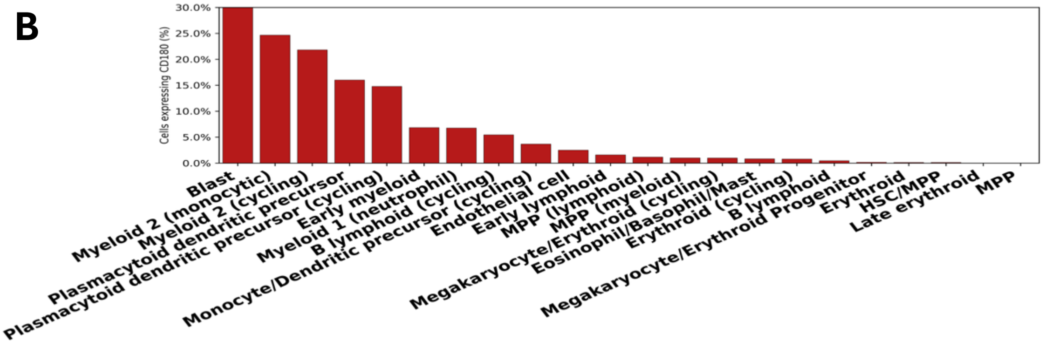

### Supplementary Figure 4

A

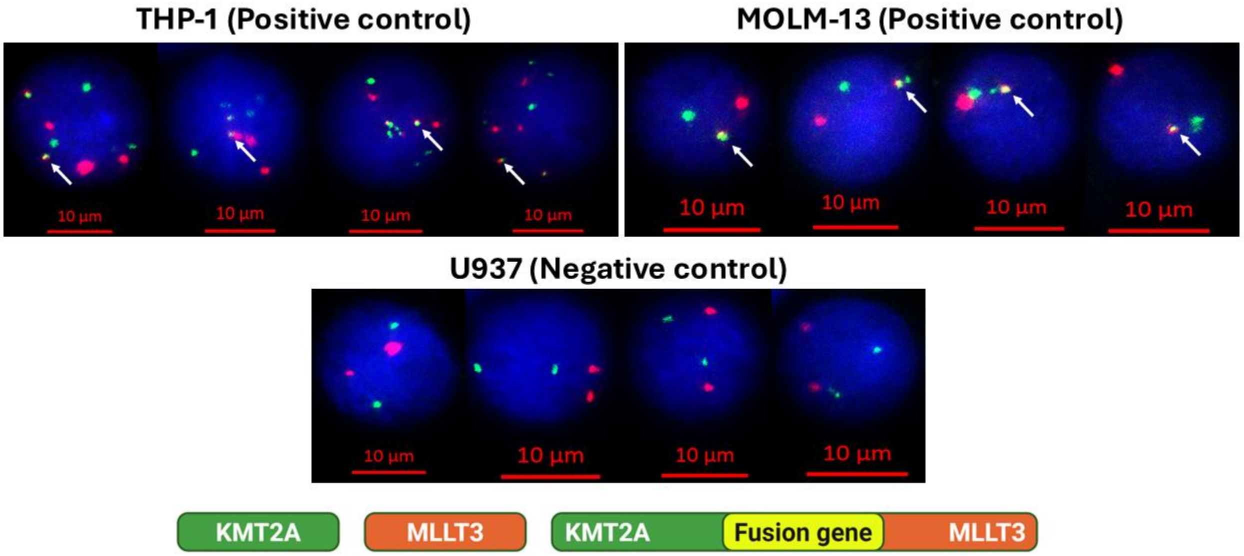


B

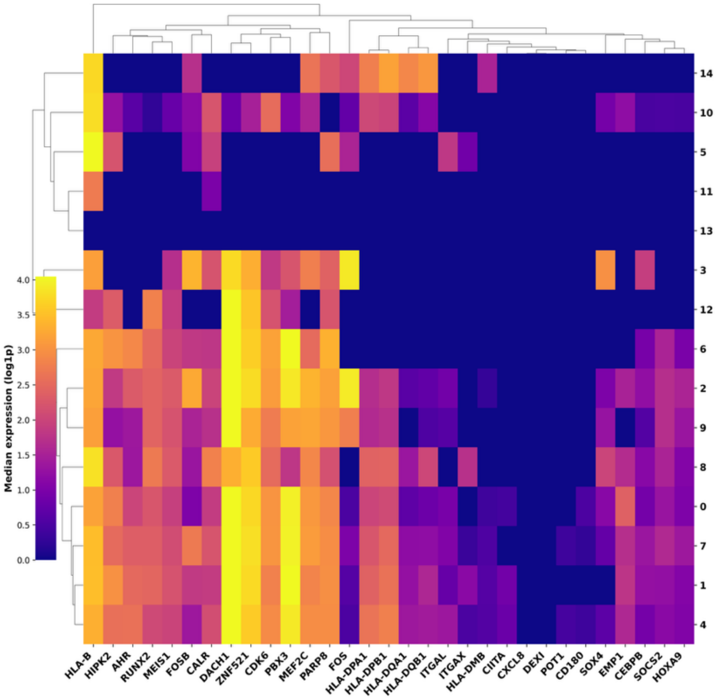

### Supplementary Figure 5

A

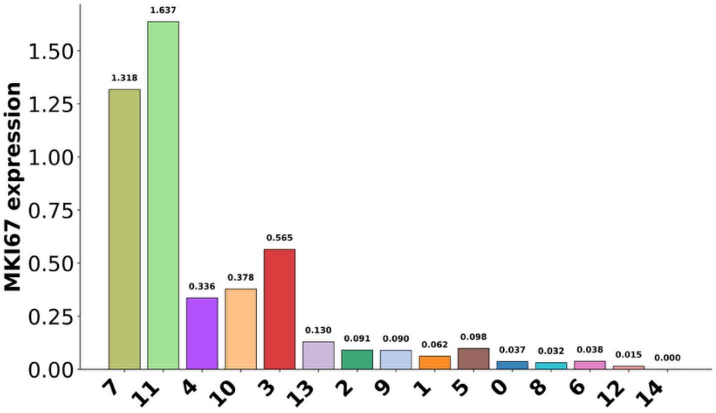


B

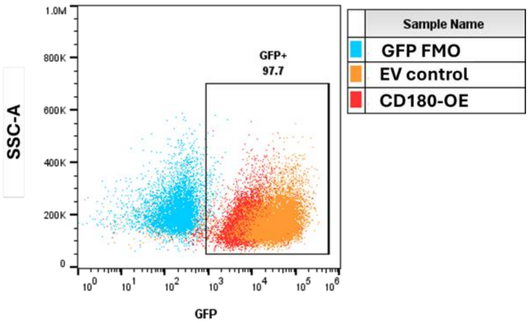

### Supplementary Figure 6

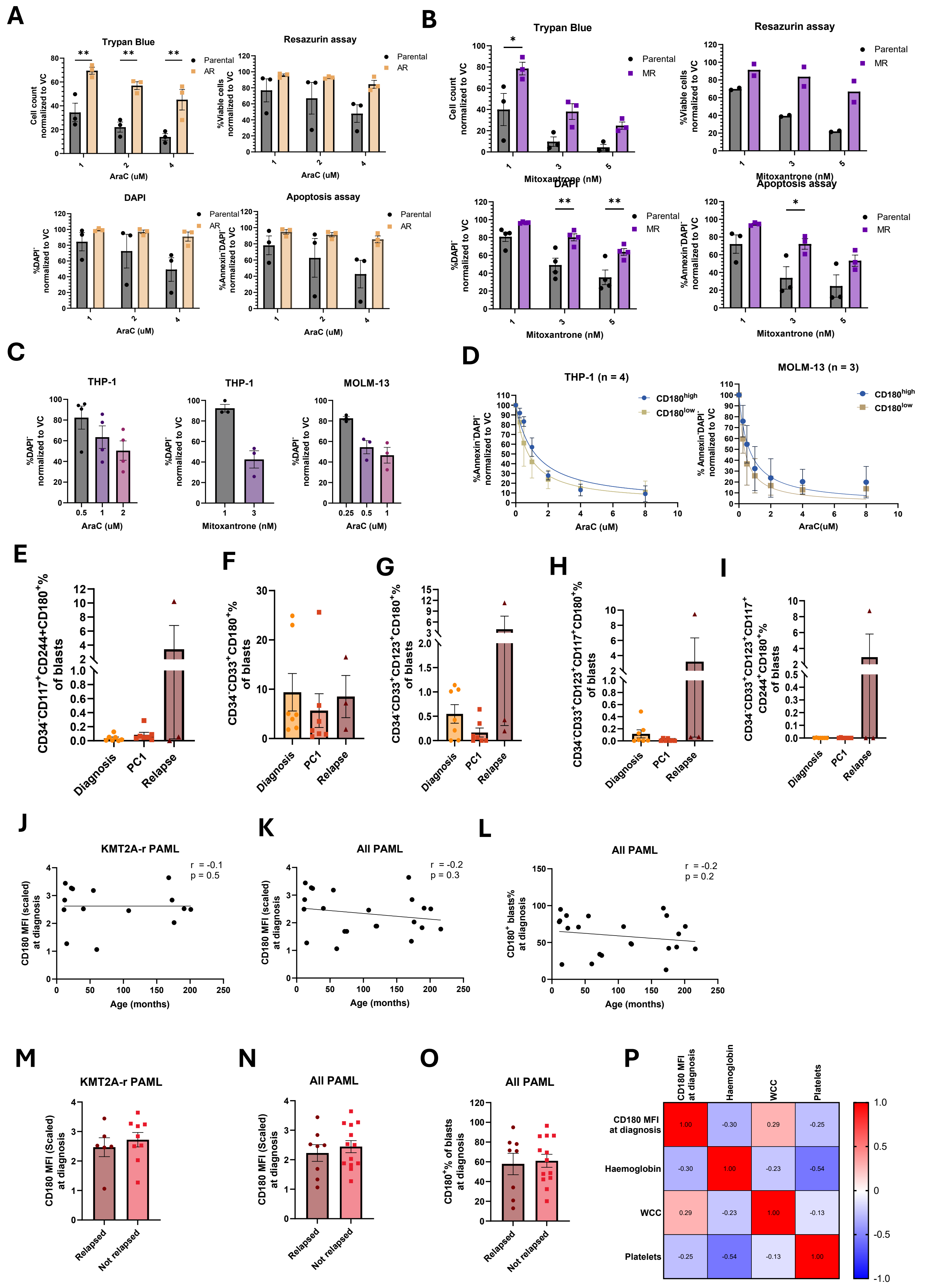

### Supplementary Figure 7

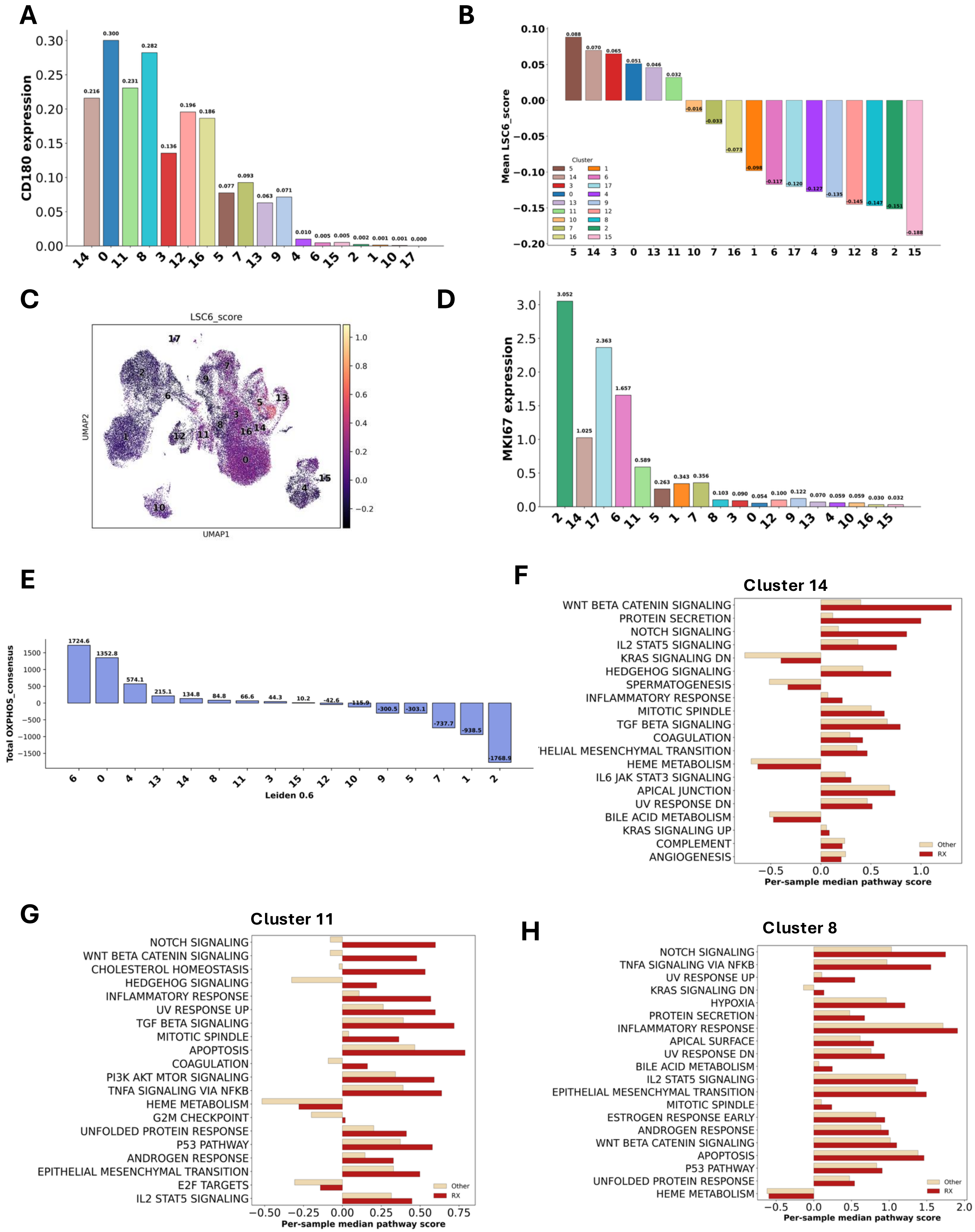
