## Supplementary figure legends for "CD180 identifies chemoresistant stem-like blasts and reveals a KMT2A-driven vulnerability in acute myeloid leukaemia"

**Supplemental Figure legends**

**Supplementary Figure 1**

**A-B)** Survival curves of the upper and lower quartile of *CD180* gene expression from *RUNX1::RUNX1T1* (t(8;21)) and *CBFB::MYH11* (inv(16)) patients respectively, with p values highlighted. B) Gating strategy for patient samples lin- CD45^low^ blast population. **C-E)** Bar plots depicting the mean arcsinh-transformed fluorescence intensity (scaled MFI; left) and the percentage of positive cells (right) for CD56, CD93, and CD36 in blasts from KMT2A-r PAML, non-KMT2A PAML, adult AML, normal CD34+ cells, and HSCs. n = 14 (CD56) and 7 (CD93, CD36) for KMT2A-r PAML; 6 (CD56) and 3 (CD93, CD36) for non-KMT2A PAML; 3 (CD56) and 1 (CD93, CD36) for adult AML; and 3 for all markers in normal CD34+ and HSCs. One-way ANOVA performed. **F)** Correlation matrix of marker expression in diagnostic KMT2A-r PAML blasts, shown as the percentage of positive cells. Pearson correlation coefficients are indicated, with red and blue shading representing strong positive and negative correlations, respectively.

**Supplementary Figure 2**

**A)** Bar plot summarising the mean gene expression (top) of *CD180* and *CD34* and highlighting the percentage of cells expressing these genes (bottom) within the whole dataset. **B)** Bar plot showing the mean *CD180* expression per cluster. **C-F)** UMAPs **C and E)** and bar graphs **D and F)** of LSC6 and PC2_34 mean scores respectively for each cluster coloured according to the integrated UMAP.**G-H)** Stacked bar plot showing output of GSEA results for cluster 4 vs all **G)** and cluster 7 vs **H).** Pathways are coloured according to their significance. **I-J)** Bar plots showing percentage engraftment of CD180^high^ and CD180^low^ sorted THP-1 cells in recipient mice 30 days post-transplantation, represented as the percentage of human CD45⁺ cells in the bone marrow (BM) I**)** and brain **J),** and the percentage of CD180+CD45+ cells in the BM (*n* = 5 per group)**K).**

**Supplementary Figure 3**

**A-B)** Bar plot showing the mean *CD180* expression A) and the percentage of *CD180+* expressing cells **B)** comparing KMT2A-r PAML blasts and all healthy PBM cell types from publically available sc-RNA-seq datasets.

**Supplemental Figure 4**

**A)** Representative Fluorescence in situ hybridization (FISH) images from control cell lines. Positive controls: THP-1 (top left) and MOLM-13 (top right); negative control: U937 (bottom). The KMT2A::MLLT3 translocation signal is shown in yellow (indicated by white arrows), along with distinct KMT2A (green) and MLLT3 (orange) signals. Nuclei are counterstained with DAPI. Images were acquired at 63× magnification, and scale bars are shown. **B)** Heatmap showing the median expression of the KMT2A gene signature genes across the clusters. Heatmap showing the median expression of all the KMT2A genes in the KMT2A score, and *CD180,* across the clusters. Log transformed gene expression values shown (high is yellow, low is purple).

**Supplemental Figure 5**

**A)** Bar plot showing the mean expression of *MKI67* expressing cells in the 4 diagnosis KMT2A::MLLT3 clustered populations**. B)** Representative FACS dot plot showing GFP+ CD180-OE and EV control THP-1 cells.

**Supplementary Figure 6**

**A-B)** Validation of AraC and mitoxantrone-resistant isogenic THP-1 cells. Bar plots show cell viability measured by trypan blue counting, resazurin assay, DAPI, and Annexin V^–^/DAPI^–^ viable cells (apoptosis assay) in AraC-resistant (AR; orange) versus parental (grey) THP-1 cells treated with 1, 2, and 4 uM AraC **A)**, and in mitoxantrone-resistant (MR; purple) versus parental (grey) THP-1 cells treated with 1, 3, and 5 nM mitoxantrone **B).** VC: Vehicle control. Two-way ANOVA was used for statistical analysis. **C)** Bar plots depicting the percentage of viable cells (DAPI^-^) of THP-1 cells treated with 0.5, 1, and 2 uM AraC (left, n = 4), THP-1 treated with 1 and 3 nM mitoxantrone (middle, n = 3), and MOLM-13 treated with 0.25, 0.5 and 1 uM AraC (right, n = 3) for 48 hours. **D)** AraC dose-response curves of CD180^high^ and CD180^low^ sorted THP-1 (n = 4, left) and MOLM-13 (n = 3, right) cells. EC50 values were calculated using non-linear regression analysis. **E-I)** Bar plots depicting the percentage representation of various potential LSC/LRC populations at different disease stages (diagnosis (n=7), post–course 1 (PC1, n=7), and relapse (n=3)) of KMT2A-r PAML. **J)** Correlation between CD180 expression in diagnostic blasts from KMT2A-r PAML, expressed as arcsinh-transformed scaled mean fluorescence intensity (scaled MFI) (n = 15), and patient age (months). **K-L)** Correlations between CD180 expression at diagnosis across the PAML cohort (n = 22) and patient age (months), shown as **K)** scaled MFI and **L)** percentage of CD180⁺ blasts. In **J-L),** scatter plots include fitted linear regression lines with Pearson correlation coefficient (*r*) and *p*-value indicated. **M)** Bar plot showing the CD180 scaled MFI of blasts at diagnosis in KMT2A-r PAML patients who relapsed (n = 6) and in patients who did not relapse by the time of analysis (n = 9). **N-O)** bar plots showing CD180 expression at diagnosis across the PAML cohort (n = 22) in patients who relapsed (n = 8) and in patients who did not relapse by the time of analysis (n = 13), shown as **N)** scaled MFI and **O)** percentage of CD180^+^ blasts. **P)** Correlation matrix of the CD180 scaled MFI of blasts at the diagnosis in KMTA-r PAML with patient haematological parameters including haemoglobin, white cell count (WCC), and platelets (n = 10). Pearson correlation coefficients are indicated, with red and blue shading representing strong positive and negative correlations, respectively.

**Supplementary Figure 7:**

**A)** Bar graph showing the mean expression of *MKI67* across all clusters in the integrated UMAP. **B-C)** Bar graphs **B)** and UMAPs **C)** of LSC6 mean score for each cluster coloured according to the integrated UMAP. **D)** Bar graph showing the mean expression of *CD180* across all clusters in the integrated UMAP. **E)** Bar plot showing total consensus score for the oxidative phosphorylation pathway activity score from each cluster in a UMAP from relapse only. **F-H)** Bar plots showing the per sample median for the combined pathway activity score for relapse cluster 14, 11 and 8 vs other disease stage cluster 14, 11 and 8.
