## supplementary table 2 for "CD180 identifies chemoresistant stem-like blasts and reveals a KMT2A-driven vulnerability in acute myeloid leukaemia"

### Table 2: Leukaemic stem cell (LSC) markers in adult versus paediatric AML.

| LSC marker | Adult AML reference and validation status | PAML reference and validation status |
|---|---|---|
| **CD34+/CD38-** | Validated as an LSC-enriched compartment (Bonnet and Dick, 1997) (Eppert et al., 2011). | Validated as an LSC-enriched compartment (Witte et al., 2011). |
| **CD123 (IL3RA)** | Validated as an LSC marker and therapeutic target ((Jin et al., 2009) (Roug et al., 2014) (Al-Mawali et al., 2017). | Less reliably validated for LSCs compared to adults. Widely expressed on PAML blasts and a potential therapeutic target. (Petersen et al., 2022) (Kandeel et al., 2021) (Naik et al., 2022). |
| **CD33** | Validated to be expressed on some LSCs while widely expressed on AML blasts. Validated as a therapeutic target (Lee et al., 2018) (Walter et al., 2012). | Validated to be expressed on some LSCs while widely expressed on PAML blasts. Validated as a therapeutic target (Pollard et al., 2021) (Willier et al., 2021). |
| **TIM3 (HAVCR2)** | Validated as an LSC marker (Saito et al., 2010) (Kikushige et al., 2015) (He et al., 2020) (Jan et al., 2011). Recently being evaluated in early-phase trials as a therapeutic target (Zeidan et al., 2024). | Less reliably validated for LSCs compared to adults. Used in PAML clinical trials for LSC and MRD detection (NCT02724163). |
| **CD47** | Validated as an LSC marker and a therapeutic target (Majeti et al., 2009) (Sick et al., 2012) (Feng et al., 2018). | Not yet validated as an LSC marker. |
| **CLEC12A (CLL-1)** | Validated as an LSC marker and a therapeutic target (Van Rhenen et al., 2007) (Lin et al., 2021). | Validated as an LSC marker and a therapeutic target (Willier et al., 2021) (Petersen et al., 2022). Used in PAML clinical trials for LSC and MRD detection (NCT02724163). |

| | | |
|---|---|---|
| **CD93** | Validated as an LSC marker and highly expressed in M4/M5 AML (Jia et al., 2022) (Iwasaki et al., 2015) (Coustan-Smith et al., 2018). | Less reliably validated for LSCs compared to adults while widely expressed on PAML blasts (Petersen et al., 2022). |
| **CD9** | Validated as an LSC marker (Liu et al., 2021) (Touzet et al., 2019). | Not yet validated as an LSC marker. |
| **PROM1 (CD133)** | Validated as an LSC marker in a subset of AML and associated with stemness and occasionally with prognosis (Heo et al., 2020) (Tolba et al., 2013). | Less reliably validated for LSCs compared to adults. Comparable expression on PAML LSCs and HSCs was detected (Cheng et al., 2016). Expression on LSCs in paediatric ALL was noticed (Cox et al., 2009). |
| **CD36** | Less reliably validated compared to other markers. May mark a subpopulation with stem-like properties (Sachs et al., 2019). | Not yet validated as an LSC marker. Limited evidence of high expression at relapse (Hoch et al., 2021). |
| **CD44** | Validated as an LSC marker and involved in LSC maintenance, adhesion, survival, and drug resistance (Jin et al., 2006) (Florian et al., 2006) (Ponta et al., 2003). | Less reliably validated for LSCs compared to adults (van der Werf et al., 2023) (Holm et al., 2015). |
| **CD56 (NCAM1)** | Validated as an aberrant MRD marker that is associated with poor prognosis (Schuurhuis et al., 2018) (Raspadori et al., 2001). | Validated as an aberrant marker helping in diagnosis and MRD tracking (Quessada et al., 2021) (Liang et al., 2022) |
| **CD117 (KIT)** | Validated as a marker of LSC-enriched population in combination with other markers in both CD34+ and CD34- AML while being expressed in normal haematopoiesis (Quek et al., 2016) (Roug et al., 2014) (Maillard et al., 2020). | Less reliably validated for LSCs compared to adults. Similar expression levels in PAML and normal haematopoietic populations (Chávez-González et al., 2014). Less established link with prognosis (Pollard et al., 2010) |

| | | |
|---|---|---|
| **MYC** | Validated to have a well-established link with LSC maintenance, metabolism, and survival programs (Amaya et al., 2022) (Nathan et al., 2024). | Not yet validated as an LSC marker. Some evidence of frequent mutation in PAML (Bolouri et al., 2017). |
| **CD244** | Validated as an LSC marker particularly of value in CD34- AML. However, it is abundantly expressed in AML at diagnosis and relapse (Haubner et al., 2018) (Quek et al., 2016) (F. Zhang et al., 2017). | Less reliably validated for LSCs compared to adults. Some evidence is present for its microenvironment role in PAML (Yuan et al., 2024) (Y. Zhang et al., 2025). |

"Validated" status indicates that the marker has been functionally or prognostically demonstrated to be associated with LSCs in primary research, while "Less reliably validated" reflects areas of ongoing research, heterogeneity, or limited specific applicability as a primary LSC-defining marker, especially when comparing to adult AML. Specific references are provided to support each entry.
