## supplementary table 3 for "CD180 identifies chemoresistant stem-like blasts and reveals a KMT2A-driven vulnerability in acute myeloid leukaemia"

### Table 3: KMT2A target genes

| Gene | Reference |
|---|---|
| *AHR* | (Fleischmann et al., 2014) *KMT2A-MLLT3* targets |
| *DRD5* | |
| *TAS1R3* | |
| *ROR2* | |
| *HIPK2* | |
| *PARP8* | |
| *ATP2B2* | |
| *EMP1* | |
| *ITGAL* | |
| *ITGAX* | |
| *CCL2* | |
| *IL8* | |
| *S100A12* | |
| *SOCS2* | |
| *CEBPB* | |
| *CIITA* | |
| *DACH1* | |
| *EGR2* | |
| *FOS* | |
| *FOSB* | |
| *HOXA11* | |
| *HOXA9* | |
| *HOXC4* | |
| *IKZF2* | |
| *MEF2C* | |
| *PBX3* | |
| *SOX4* | |
| *SP9* | |
| *ZNF521* | |
| *CALR* | |
| *CGNL1* | |
| *DEXI* | |
| *GPNMB* | |
| *HLA-B* | |
| *HLA-DMB* | |
| *HLA-DPA1* | |
| *HLA-DPB1* | |
| *HLA-DQA1* | |
| *HLA-DQB1* | |
| *HLA-DRB4* | |
| *POT1* | |
| ***MEIS1*** | *Essential KMT2A*-r targets (Grembecka et al., 2012) (Placke et al., 2014) (Kuo et al., 2009) |
| ***CDK6*** | |
| ***RUNX2*** | |

Fleischmann, K. K., Pagel, P., Schmid, I., & Roscher, A. A. (2014). RNAi-mediated silencing of MLL-AF9 reveals leukemia-associated downstream targets and processes. *Molecular Cancer*, *13*(1), 1–14. https://doi.org/10.1186/1476-4598-13-27/FIGURES/7

Grembecka, jolanta, He, shihan, shi, aibin, purohit, trupta, Muntean, andrew G., sorenson, roderick, showalter, H., Murai, M., Belcher, amalia M., Hartley, thomas, Hess, jay L., & Cierpicki, tomasz. (2012). Menin-MLL inhibitors reverse oncogenic activity of MLL fusion proteins in leukemia. *Nature CHeMICaL BIOLOGY |*, *8*. https://doi.org/10.1038/nCHeMBIO.773

Kuo, Y. H., Zaidi, S. K., Gornostaeva, S., Komori, T., Stein, G. S., & Castilla, L. H. (2009). Runx2 induces acute myeloid leukemia in cooperation with Cbfβ-SMMHC in mice. *Blood*, *113*(14), 3323. https://doi.org/10.1182/BLOOD-2008-06-162248

Placke, T., Faber, K., Nonami, A., Putwain, S. L., Salih, H. R., Heidel, F. H., Krämer, A., Root, D. E., Barbie, D. A., Krivtsov, A. V., Armstrong, S. A., Hahn, W. C., Huntly, B. J., Sykes, S. M., Milsom, M. D., Scholl, C., & Fröhling, S. (2014). Requirement for CDK6 in MLL-rearranged acute myeloid leukemia. *Blood*, *124*(1), 13–23. https://doi.org/10.1182/BLOOD-2014-02-558114,
