## Supplementary Materials and Methods for "CD180 identifies chemoresistant stem-like blasts and reveals a KMT2A-driven vulnerability in acute myeloid leukaemia"

**Single-cell RNA-sequencing**

Single-cell RNA sequencing libraries were generated using the Chromium Next GEM Single Cell 3′ Reagent Kits v3.1 (10x Genomics, Pleasanton, CA, USA). Primary patient samples were sorted into the required cell populations and prepared at 700 cells/µl according to the manufacturer’s instructions. GEM generation, barcoding, reverse transcription, cDNA amplification, and library construction were performed following the Chromium Next GEM Single Cell 3′ Gene Expression v3.1 Dual Index protocol (10x Genomics). For GEM generation, 23.6 µl of cell suspension was combined with nuclease-free water, the GEM generation master mix, and partitioning oil on a Chromium Next GEM Chip G, and processed using the Chromium Controller. Single cells were encapsulated at limiting dilution to ensure that most GEMs contained either zero or one cell. Upon GEM formation, barcoded gel beads dissolved to release primers consisting of the TruSeq Read 1 sequence, a 16-nt 10x cell barcode, a 12-nt UMI, and a poly(dT) sequence for mRNA capture. Reverse transcription occurred within each GEM, producing barcoded cDNA from polyadenylated transcripts. Following GEM-RT, cDNA was recovered by Silane bead purification, eluted, and PCR-amplified to generate sufficient material for library construction. 3′ gene expression libraries were then prepared via enzymatic fragmentation, size selection, adaptor ligation, and a final PCR amplification. Library size distributions (target range 200–900 bp) and cDNA quality were assessed using an Agilent Bioanalyzer. Sequencing was performed at Glasgow Polyomics on an Illumina NovaSeq platform using the recommended cycle configuration: 28 cycles for Read 1 (cell barcode, UMI, and 5′ transcript end), 10 cycles for index reads, and 90 cycles for Read 2 (3′ transcript end). Each sample was sequenced to a depth of approximately 400 million reads.

**Single cell data analysis**

**Processing & quantification.** FASTQs were processed using Cell Ranger on default settings again GRCh38. The outputs of which were imported into Scanpy (1.10.4). **Quality control.** Cells were filtered using Scanpy with the use of thresholds being defined as n MADS from the median for n_genes, n_counts and % mitochondrial. Alongside this Ambient RNA correction was done using SoupX with default parameters. Doublets were assessed with Scrublet (0.2.3) but provided negligable changes, so unfiltered data was used thereafter. **Normalisation, features, and integration.** Data were normalized and log transformed using Scanpy; highly variable genes were selected using scanpy.pp.highly_variable_genes. The top 4000 genes were selected and used for the cross-dataset integration using scVI(scvi-tools 1.3.0), when integrating across timepoints the timepoints were used as categorical covariates during model training. Default hyperparameteers were used unless stated otherwise. Once integrated the full selection of genes were added back to the integrated object. **Dimensionality reduction and clustering.** PCA was performed then followed by UMAP. Leiden Clustering was performed, and the optimal resolution was determined by clustering over a grid of resolutions and jointly maximising silhouette score and Rand-index-based stability metric across random seeds. **Differential expression and pathway analysis.** Differential expression was performed using Scanpy where multiple-testing correction by Benjamini–Hochberg (q < 0.05) was performed. Pathway activity per cell/sample was measured using a combination of decoupler’s (1.9.2) ULM and AUCell scores that were combined into a z-score composite. Any pathway activity or GSEA used only MSigDB Hallmark collection. Additional gene scores were computed using Scanpy’s score_genes. **Visualization & survival.** All plots were generated in Python; survival curves were produced in R using the packages survival and survminer.

**ATAC-sequencing**

AML cells were harvested and washed twice with ice cold phosphate buffered saline (PBS) and counted to ensure high viability (>90%), 50,000 viable cells per sample. Cells were centrifuged at 300g for 5 minutes in 4◦C. Cell pellets were resuspended in 50 µL of ice-cold lysis buffer consisting of 10 mM Tris-HCl (pH 7.4), 10 mM NaCl, 3 mM MgCl₂, 0.1% IGEPAL CA-630, 0.1% Tween-20, and 0.01% digitonin and incubated on ice for 3 minutes, followed by addition of 1 mL of wash buffer (10 mM Tris-HCl, 10 mM NaCl, 3 mM MgCl₂, 0.1% Tween-20) to remove excess detergents. Nuclei were pelleted by centrifugation at 500g for 10 minutes at 4°C and resuspended in a 50 µL transposition reaction mix containing 25 µL 2× TD buffer, 2.5 µL Tn5 transposase (Illumina Nextera), and 22.5 µL nuclease-free water. The transposition reaction was incubated at 37°C for 30 minutes in a thermomixer with gentle shaking to ensure uniform enzyme accessibility. Transposed DNA fragments were purified using a MinElute PCR Purification Kit (Qiagen) according to the manufacturer’s instructions and eluted in 10 µL of elution buffer. Sequencing libraries were prepared by PCR amplification of the transposed DNA using indexed primers (25 μM each). Amplification was performed in a thermocycler under the following conditions: an initial extension at 72°C for 5 minutes, denaturation at 98°C for 30 seconds, followed by 10 amplification cycles consisting of 98°C for 10 seconds, 63°C for 30 seconds, and 72°C for 1 minute. Amplified libraries were purified using 1.2x SPRI magnetic bead selection to select the right sized DNA fragments. The beads were washed twice with 180μL of 80% ethanol, and the DNA was eluted in 25μL of elution buffer. Quality control (QC) and library size distribution were assessed using the Agilent Bioanalyzer. DNA concentration was measured with the Qubit High-Sensitivity DNA Assay Kit. High quality libraries was confirmed by having the characteristic nucleosomal ladder pattern, corresponding to fragments around 200 bp (nucleosome-free), 400 bp (mononucleosome), and 600 bp (dinucleosome). After QC confirmation, libraries were stored in Lo-Bind tubes at -20 ˚C for sequencing. The libraries were sent to Novogene for sequencing. Sequencing was performed using paired end reads (2 × 150 bp) on an Illumina NovaSeq X Plus with sequencing depth of 50 million reads per sample to achieve comprehensive coverage of accessible chromatin regions.

**Bulk RNA-sequencing**

RNA was harvested using either the RNeasy Mini Kit (Qiagen) or Arcturus PicoPure kit (Thermofisher Scientific). Thereafter, before RNA sequencing, RNA quality was checked for all the samples using Bioanalyzer. The polyA RNA libraries were prepared using the TruSeq stranded mRNA kit (Illumina). Paired end sequencing was generated on NextSeq2000 that yields a read of 100 bp to a depth of 30 million reads.

**ChIP-sequencing**

Chromatin immunoprecipitation was conducted as previously described (42). Briefly, 10^7^ cells were single-fixed (1% formaldehyde for 10 min) for histone modifications or double-fixed (2 mM disuccinimidyl glutarate for 30 min, then 1% formaldehyde for 30 min) for transcription/chromatin factors, then lysed with 120 µl SDS lysis buffer (10 mM Tris-HCl pH 8.0, 1 mM EDTA, 1% SDS) and sonicated using a Covaris ME220 (Woburn, MA) to generate 200–300 bp fragments. Insoluble material was pelleted, then the supernatant was diluted 10× and pre-cleared for 30 min at 4 °C with rotation using 5 µl protein A- and G-coupled dynabeads (ThermoFisher Scientific). An input sample was taken, then 2 µg antibody was added to the sample before incubation overnight at 4 °C with rotation. The antibodies used were: H3K4me1 (pAb-194-050, diagenode), H3K4me3 (39159, active motif), H3K27ac (C15410196, diagenode), H3K79me3 (C15410068, diagenode), KMT2A (A300-086A, Bethyl) and BRD4 (A301-985A lot 1, Bethyl). Protein A- and G-coupled dynabeads were used to isolate antibody-chromatin complexes, after which the beads were washed three times with RIPA buffer (50 mM HEPES-KOH pH 7.6, 500 mM LiCl, 1 mM EDTA, 1% NP40 and 0.7% sodium deoxycholate) and once with Tris-EDTA. Samples were eluted with SDS lysis buffer, RNase A- and proteinase K-treated and crosslinks were reversed at 65 °C overnight. DNA was purified by PCR purification column (Qiagen) and analyzed by qPCR, relative to input. For ChIP-seq, libraries were generated using the Ultra II library preparation kit (NEB), then sequenced by paired-end sequencing with a 75-cycle high-output Nextseq 500 kit (Illumina).

**Epigenomic data analysis**

ChIP-seq, ATAC-seq and bulk RNA-seq analysis was performed using the SeqNado pipeline (https://github.com/alsmith151/SeqNado). In brief, following QC of FASTQ files by FastQC (v0.12.1;4), reads were trimmed using trim_galore with Cutadapt (v0.6.10). Trimmed reads were then mapped to the hg38 reference genome using Bowtie 2 (v2.5.1) or STAR (v2.7.11b) for RNA-seq samples. PCR duplicates were removed for ChIP-seq and ATAC-seq samples using picard MarkDuplicates (v3.0.0). Problematic genomic regions present in the ENCODE Blacklist were removed from the aligned files, and further QC of the aligned files was performed using samtools (v1.17). BigWigs were generated using the deepTools (v3.5.1) bamCoverage command (https://github.com/deeptools/deepTools), with the flags extendReads –normalizeUsing RPKM. For RNA-seq counts from the forward and reverse strand were separated using --filterRNAstrand [forward|reverse] and counts for the reverse inverted for visualisation using --scaleFactor -1. Visualization was performed in the UCSC genome browser https://genome-euro.ucsc.edu/s/asmith151/2025%2D09%2D17%2DTHP1%2Dsession. ChIP-seq datasets include GSE117865, GSM2212253, GSM3983886, GSM2108045.

**Computational Data analysis**

Gene expression data were retrieved and visualized using BloodSpot 3.0 to compare selected genes across normal and malignant haematopoietic populations (Bloodpool:AML samples with normal cells). All data were accessed using the default normalization settings provided by the platform.

**Tissue Culture**

All tissue culture procedures were conducted under aseptic conditions using Class I biosafety cabinets for established cell lines and Class II cabinets for primary patient samples. Cells were maintained at 37 °C in a humidified incubator with 5% CO₂. Human cell lines were obtained from a certified commercial source (ATCC®, TIB-202 for THP-1 and DSMZ, ACC 554 for MOLM-13) and cultured according to the manufacturer’s recommendations and standard mammalian cell culture protocols. Routine mycoplasma testing was performed using qPCR-based detection kits (Venor®GeM qOneStep or MycoAlert), and all cell lines were confirmed to be mycoplasma-free and authenticated. Adherent HEK293T cells (ATCC®, CRL-3216) were passaged using 0.25% trypsin–EDTA, neutralized with complete medium, and reseeded as required.

**Drug treatment and Cell Viability assays**

Cytarabine (AraC; MW = 243.22 g/mol) and mitoxantrone (MW = 517.4 g/mol) were obtained as powders (Sigma #C3350000 and Selleckchem #S2485, respectively) and reconstituted in distilled water or DMSO, respectively, according to the manufacturer’s instructions. Aliquots were stored under recommended conditions until use. For treatment experiments, cells were seeded at a density of 0.2 × 10⁶ cells/mL and treated 24 hours later with freshly prepared drug dilutions in complete medium. Treatments were performed for 48–72 hours, depending on experimental design. Vehicle controls were included for each drug (media-only for AraC and 0.01% DMSO for mitoxantrone). Dose–response assays were performed in THP-1 and MOLM-13 cells to determine the half-maximal effective concentration (EC₅₀) of AraC and mitoxantrone. Cells were treated with increasing drug concentrations alongside untreated and vehicle controls. Cell viability was assessed using the trypan blue (Gibco #15250061) exclusion assay, resazurin assay (Merck #R7017) for evaluating cell metabolic activity, and assessing apoptotic status using Annexin V (Biolegend #640945) and DAPI (Oxoid #BR014G) staining. Apoptosis assay was performed by harvesting between 0.05–0.5 × 10⁶, washing with Hanks’ buffered saline solution, and staining with a master mix of Annexin V and DAPI for 15 minutes at room temperature in the dark.

**Apoptosis Assay**

Apoptosis was assessed using Annexin V/DAPI staining. Following drug treatment, 0.05–0.5 × 10⁶ cells were harvested, washed in HBSS, and incubated with Annexin V and DAPI for 15 min at room temperature in the dark. Samples were diluted with HBSS and analysed by flow cytometry. Viable cells were defined as Annexin V⁻/DAPI⁻. Appropriate single-stain controls were included to confirm gating. Data were analysed using FlowJo v10 and GraphPad Prism 9.

**Cell Cycle Assay**

Cell cycle distribution was assessed in sorted cell lines and patient samples using Ki67 (Biolegend #350526) and DAPI staining. Cells were allowed to recover for 24 hours post-sorting, then fixed and permeabilized using the Cytofix/Cytoperm™ kit (BD Biosciences #554714) according to the manufacturer’s instructions. After fixation, cells were incubated with Ki67 antibody at 4 °C for 30 minutes, followed by DAPI staining for 10 minutes at room temperature in the dark. Flow cytometry was used to analyze DAPI and Ki67 fluorescence. Doublets and apoptotic cells with very low DNA content were excluded to ensure accurate assignment of cell cycle stages. G0 (quiescent), G1, S, G2, and M phases were determined based on combined DAPI and Ki67 staining patterns.

**Primary Patient Samples**

Primary bone marrow samples were obtained from the Myechild01 clinical trial, the VIVO Biobank (UK), the Haematology Biobank at the University of Glasgow, and the Schehallion Ward at Queen Elizabeth University Hospital (QEUH), Glasgow. All patients or guardians provided written informed consent, and ethical approval was obtained in accordance with West of Scotland Research Ethics protocols (adult and paediatric, reference 20-WS-0066). Samples from the Myechild01 trial and VIVO Biobank were collected under material transfer agreements with the University of Birmingham, University of Oxford and the University of Newcastle. Fresh samples received directly from hospitals were processed on the day of collection to isolate mononuclear cells (MNCs) for cryopreservation, while other samples were supplied as pre-processed, frozen MNC cryovials. Cord blood–derived mononuclear cells (CBDMC) were purchased from STEMCELL Technologies (70007.1).

**Isolation and Recovery of Mononuclear Cells**

Bone marrow (BM) samples requiring MNC isolation were collected in EDTA-containing tubes and filtered through a 100 µm strainer to remove clots and debris. Cells were centrifuged to remove plasma. The cell pellet was resuspended in buffer and layered over Histopaque for density gradient centrifugation. MNCs were collected from the interface, washed, counted, and either used fresh or cryopreserved in freezing medium at a 1:1 ratio. Cryovials were initially stored at −80 °C in a controlled-rate freezing container before transfer to liquid nitrogen for long-term storage. For recovery, cryopreserved MNCs were thawed in a 37 °C water bath and gradually diluted with thawing medium. Cells were washed, filtered through a 100 µm strainer, counted, and cultured at 1–2 × 10⁶ cells/mL for 2 hours prior to downstream applications.

**Flow** **cytometry**

**Panel design:** We designed an in-house 19-marker flow cytometry panel to comprehensively characterize leukaemic stem cell (LSC) populations while simultaneously excluding lymphoid cells. The panel included the following markers with their corresponding fluorophores, suppliers, and catalogue numbers: CD34 (BV421, BD Biosciences #562577), CD33 (BV510, BD Biosciences #563257), CD117 (BV650, BD Biosciences #563859), CD56 (BV711, BD Biosciences #563169), CD45RA (BV786, BD Biosciences #741010), CD38 (BB515, BD Biosciences #564498), CD244 (PerCP-Cy5, BioLegend #329516), CD180 (PE, BioLegend #312906), CD90 (PE-Cy7, BD Biosciences #561558), CD123 (R718, BD Biosciences #567287), CD45 (APC-Cy7, BD Biosciences #557833), 7-AAD (7-AAD, BD Biosciences #559925), CD3 (PE-Cy5, BD Biosciences #555341), CD4 (PE-Cy5, BD Biosciences #555348), CD8 (PE-Cy5, BD Biosciences #555636), CD19 (PE-Cy5, BD Biosciences #560993), CD20 (PE-Cy5, BD Biosciences #555624), CD36 (BV605, BD Biosciences #563518), and CD93 (APC, BioLegend #336120). Panel design considered fluorophore brightness relative to expected antigen expression, minimized spectral overlap between co-expressed markers (e.g., CD123 and CD45RA on GMPs), and accounted for instrument configuration to maximize resolution of rare subpopulations.

**Flow Cytometry Staining and Sorting**

For cell lines, 0.2–0.5 × 10⁶ cells per sample were harvested, washed with PBS containing 2% FBS, and incubated with fluorochrome-conjugated antibodies for 30 min on ice. Cells were washed and resuspended in PBS/2% FBS containing a viability dye (e.g., DAPI) prior to data acquisition. For primary patient samples, cells were harvested and counted, then, to reduce non-specific binding in multiparametric panels, Fc receptor blocking reagent (BD Biosciences #564220) and BD Brilliant Stain Buffer (BD Biosciences #563794) were added and incubated for 10 min before antibody addition. Antibody master mixes were prepared in PBS/2% FBS, added to samples, and incubated for ≥30 min on ice. Cells were then washed, resuspended in PBS/2% FBS with 7-AAD, and analysed. Flow Cytometry Analysis was performed on the Sony ID7000 and Fluorescence-activated cell sorting (FACS) was performed on a BD FACSAria II (BD Biosciences) using FACSDiva software. CD180+ and CD180- populations were isolated from cell lines and primary samples for downstream assays. Unstained and FMO controls were used to define gates. Primary cells were rested for 2 h post-thaw prior to staining and sorting to allow recovery while minimizing culture-induced phenotypic changes, particularly in stem cell populations.

**Flow Cytometry Data Analysis**

Flow cytometry data were analysed using FlowJo v10 (BD Biosciences) and the OMIQ cloud-based platform. FlowJo was used for standard analyses employing manual gating. For high-dimensional, multiparametric datasets, OMIQ was used due to its integrated tools for batch correction, automated clustering (e.g., FlowSOM), and dimensionality reduction (e.g., UMAP), enabling efficient large-scale data processing. Surface marker expression was reported as either geometric mean fluorescence intensity (MFI) normalized to the appropriate IgG or FMO control, or as mean arcsinh-transformed fluorescence intensity (scaled MFI). Arcsinh transformation (cofactor = 400) was used to ensure comparability across samples by compressing high-intensity signals while preserving resolution of lower-intensity populations. Mean arcsinh-transformed fluorescence intensities (“scaled MFI”) were exported for each population.

**Fluorescence In Situ Hybridization (FISH)**

FISH was performed to detect the KMT2A-MLLT3 t(9;11) translocation in sorted CD180+ and CD180- primary patient cells. A dual-fusion probe was used (MetaSystems #D-5133-100-OG), consisting of a green-labelled KMT2A probe (11q23.3) and an orange-labelled MLLT3 probe (9p21); fusion (yellow) signals indicated a rearrangement. THP-1 and MOLM-13 cells served as positive controls, and U937 cells as negative control. For each sample, 0.02–0.2 × 10⁶ cells were subjected to hypotonic treatment (0.075 M KCl, 37°C, 15 min) followed by fixation in methanol: acetic acid. Fixed cells were applied to poly-L-lysine–coated slides and air-dried. Probes were applied, and slides were denatured at 75°C for 2 min, followed by overnight hybridization at 37°C. After post-hybridization washes, nuclei were counterstained with DAPI and imaged using a Zeiss LSM 780 fluorescence microscope. At least two independent scorers evaluated signal patterns, applying predefined inclusion/exclusion criteria to ensure accurate interpretation.

**CRISPR–Cas9 Knockout of CD180**

CD180 knockout (CD180-KO) was generated in THP-1 cells using the lentiCRISPR v2 system (Addgene #52961), which expresses both Cas9 and the sgRNA. sgRNA sequences targeting CD180 were designed and the four highest-scoring guides (minimal predicted off-target activity) were cloned into lentiCRISPR v2. CD180 Sg1: forward CACCG**TCTGCTGGGACAAGATAGGG** and reverse AAAC**CCCTATCTTGTCCCAGCAGA**C; CD180 Sg2: forward CACCG**AATGCACATCTGATCCCAGG** and reverse AAAC**CCTGGGATCAGATGTGCATT**C; CD180 Sg3 forward CACCG**CGTCAGCTGCTTCTTTTGGG** and reverse AAAC**CCCAAAAGAAGCAGCTGACG**C; CD180 Sg4 forward CACCG**AAGGTTCTATTGTGAATTGT** and reverse AAAC**ACAATTCACAATAGAACCTT**C. A non-targeting sgRNA in the same vector was used as a control. Recombinant lentiviral particles were produced and used to transduce THP-1 cells, followed by antibiotic selection to establish stable knockout lines. Knockout efficiency was verified by flow cytometry.

**Generation of CD180 Knockout THP-1 Cells**

Single guide RNAs (sgRNAs) targeting CD180 were cloned into the lentiCRISPR v2 vector and packaged into lentiviral particles using HEK 293T cells co-transfected with psPAX2 and pVSVG plasmids via polyethyleneimine (PEI). Viral supernatants were collected, filtered, and used to transduce THP-1 cells in the presence of polybrene, followed by puromycin selection to enrich transduced populations. Knockout efficiency was confirmed by flow cytometry. A non-targeting sgRNA served as a control.

**Retrovirus-Mediated CD180 Overexpression**

A custom pMSCV-PIG (Puro IRES GFP) retroviral vector containing human CD180 cDNA (GenScript, Oxford, UK; #SC2049-2) was used to generate stable CD180 overexpression. Disclaimer: the pMSCV-PIG vector was originally a gift from David Bartel to Addgene (Addgene plasmid #21654; <http://n2t.net/addgene:21654>) (Mayr and Bartel, 2009). The CD180 insert was cloned upstream of the PGK promoter between the XhoI and HpaI restriction sites, enabling co-expression of GFP via an internal ribosome entry site (IRES) as a marker of successful transduction. The construct was sequence-verified prior to use. The empty pMSCV-PIG vector served as the negative control. Retroviral particles were produced by co-transfecting HEK 293T cells with three plasmids: pMSCV-PIG (CD180 or empty vector), pCGP (packaging; Addgene #51476), and pVSVG (envelope; Addgene #138479), using polyethyleneimine (PEI) as the transfection reagent. Viral supernatants were collected and used to transduce THP-1 cells. Stable integration of the transgene was achieved through retroviral reverse transcription and integration, followed by selection with puromycin. Successful transduction and CD180 overexpression were validated by assessing GFP fluorescence and CD180 surface expression via flow cytometry. Three independent biological replicates, each involving separate transfection and transduction cycles, were performed.

**Patient-Derived Xenograft (PDX) Models**

Immune deficient NOD-SCID IL2Rγnull (NSG) and NOD/Rag1/2-/- IL2Rγ-/- (NRG) mice were engrafted with 1 × 10⁶ mononuclear cells (MNCs) from acute myeloid leukemia (AML) patient samples. After 16 weeks, BM cells were harvested, and human cell engraftment was assessed by flow cytometry based on human CD45 expression. The percentages of BM engraftment were 5.2%, 24.8%, and 13% for AML03, AML04, and PAML22, respectively. Prior to cryopreservation, BM cells were enriched for human CD45+ cells using magnetic-activated cell sorting (MACS; CD45 MicroBeads, Miltenyi Biotec, UK). Both pre- and post-engraftment samples were stained and immunophenotypically profiled using our LSC panel. Blasts were identified by gating on Lin- CD45^low^ cells (**Supplemental Figure 1B**).

***In Vivo* Transplantation of THP-1 Cells**

NRG-W41 mice (NOD/Rag1/2−/− IL2Rγ−/− KitW41/W41) and NSG were used to assess the engraftment potential of CD180+ and CD180- sorted and WT and KO THP-1 cell populations. 5 mice were transplanted with 0.5 × 10⁶ sorted THP-1 per group. Mice were monitored for 30 days following transplantation and sacrificed at the experimental endpoint. Bone marrow and brain tissues were collected, and human leukemic engraftment was quantified by flow cytometric detection of human CD45+ cells. NSG (NOD.Cg-Prkdcˢᶜⁱᵈ Il2rgᵗᵐ¹ᵂʲˡ/SzJ) mice were used for tail vein transplantation of NTC and CD180 KO generated in luciferase-expressing THP-1 cells. 10 mice were transplanted with 2 × 10⁶ NTC/KO cells per group. Leukaemic burden was assessed at day 7 and day 17 using bioluminescence imaging. Mice were injected subcutaneously with 200 μL L d-Luciferin (Beetle Luciferin, Potassium Salt cat no. E1605, Promega) and anaesthetised with 2–3% isoflurane. Images were taken with the fixed exposure using IVIS Spectrum (PerkinElmer). Signal intensity was quantified using AURA software.

**Statistical analysis**

Statistical analyses and data normality check were performed using GraphPad Prism version 9.5.. For comparisons between two groups, paired or unpaired two-tailed *t*-tests were applied as appropriate, while the Mann–Whitney *U* test was used for non-parametric two-group comparisons. One-way ANOVA with Bonferroni correction was used for analyses involving a single variable across multiple groups, and two-way ANOVA for multiple independent variables. Non-parametric data were analysed using the Kruskal–Wallis test with Dunn’s post-hoc correction. Flow cytometry marker expression was presented as arcsinh-transformed mean fluorescence intensity (scaled MFI) or as geometric MFI normalized to the corresponding isotype (IgG) or fluorescence-minus-one (FMO) control. Error bars represent the standard error of the mean (SEM). Statistical significance was defined as follows: ns, not significant; * p < 0.05; ** p < 0.01; *** p < 0.001; **** p < 0.0001. Figures were generated using GraphPad Prism, Microsoft PowerPoint, and BioRender (2025), while tables and pie charts were created in Microsoft Excel.
